# A systematic imputation framework for sparse, multimodal space biology datasets: application to retinal imaging and omics from the RR9 mission

**DOI:** 10.64898/2026.06.09.730965

**Authors:** Vaishnavi Nagesh, Lauren Sanders, Sylvain V. Costes, Pinar Avci, Ayse Yigit, Amey Agarwal, Alireza Hayati, Alavia Batool, Fathi Karouia, Atul M. Chander, Caleb M. Schmidt, Jian Gong

## Abstract

Missing data is a fundamental challenge in space biology, where high experimental costs, limited sample availability, and tissue allocation constraints produce datasets that are sparse, multimodal, and heterogeneous. We present a systematic four-stage framework for diagnosing, implementing, and validating data imputation strategies tailored to these characteristics, and demonstrate its application to retinal imaging and omics data from the NASA Rodent Research 9 (RR9) mission. Using logistic regression-based missingness diagnosis, we identify a Missing At Random (MAR) mechanism driven by experimental design constraints across nine assay modalities. We implement and optimize three imputation strategies: K-Nearest Neighbors (KNN), Multiple Imputation by Chained Equations with weak ElasticNet regularization (MICE-Elastic), and a per-column hybrid strategy, evaluated against a random sample imputer baseline. Validation across seven complementary metrics including supervised classification, unsupervised clustering, correlation structure preservation, masked value recovery, cross-dataset generalization, and permutation testing reveals that MICE-Elastic and the Hybrid strategy preserve genuine biological signal in both RNA-seq and TUNEL modalities, while KNN and the random sample imputer do not despite achieving comparable cross-validation accuracy. A critical finding is that imputation substantially improves supervised classification performance while consistently degrading unsupervised clustering structure, a trade-off researchers must understand before applying these methods. This framework provides practical, actionable guidance for space biologists and data scientists managing sparse multimodal datasets, and represents a foundational step toward digital twin development for space medicine.

## 1. Introduction

The field of space biology generates critical insights into how biological systems adapt to microgravity, radiation, and other unique stressors of the space environment (1). However, due to the high cost, limited access, and logistical constraints of conducting biological experiments in space, datasets from these studies are often sparse, incomplete, and heterogeneous in structure (2). Although ground-based analogs simulate key spaceflight stressors, they remain partial representations of the in-flight environment and do not capture the combined effects of concurrent mission stressors (3). One approach to overcome the challenge of flying samples/ people to space is the development of digital twins - computational models that simulate individual physiology in response to spaceflight stressors. Digital twins enable hypothesis generation, counterfactual prediction, and personalized risk assessment (4), making them valuable tools for space medicine. Yet realizing this potential depends on data quality. Digital twin development requires complete, high-quality training datasets. The inherent sparsity of space biology data - due to limited sample availability and instrument detection limits - presents a critical barrier to digital twin training. Thus, robust imputation that preserves biological signal while enabling model training is a prerequisite for space medicine digital twins.

One foundational step toward reliable data-driven modeling is mitigating the impact of incomplete or missing measurements through effective data imputation. To enable digital twin training on space-derived datasets, imputation must preserve biological signal while handling missing data. Robust imputation methods can reconstruct incomplete measurements, integrate cross-platform data, and enable downstream tasks such as mechanistic modeling and generative machine learning (5). Despite advances in batch-effect correction (6), the efficacy of imputation strategies for sparse, heterogeneous, high-dimensional space biology datasets remains largely unevaluated.

The combination of three factors - small sample sizes (limiting statistical power), multimodal data (heterogenous characteristics), and mixed missingness patterns (violating standard assumptions) create a unique challenge space that standard imputation methods are not designed to handle. These combined challenges necessitate a systematic methodological framework tailored to this domain.

Here we present such a framework and demonstrate its utility on a high-value case study: Spaceflight-Associated Neuro-ocular Syndrome (SANS) from the NASA Rodent Research 9 (RR9) mission, which affects a substantial proportion of astronauts during long-duration missions (7) and requires integration of multiple biological modalities (e.g. imaging, transcriptomics, plasma biomarker assays) to understand mechanisms. SANS thus represents an ideal test case for our multimodal imputation framework. The retina serves as a non-invasive window into systemic physiology (8), making retinal multimodal data from the RR9 mission an ideal testbed.

Our framework:

1. Systematically diagnoses missingness mechanisms.
2. Implements and optimizes multiple imputation strategies.
3. Evaluates their impact on downstream analytical tasks, providing practical guidance for researchers managing heterogeneous, multimodal, sparse datasets in space biology

## 2. Author Summary

Space biology experiments are expensive, logistically complex, and inherently limited in sample size, resulting in datasets that are frequently incomplete and highly heterogeneous (2). Missing data is a fundamental barrier to building reliable computational models of how the human body responds to spaceflight. This work introduces a systematic framework for addressing missing data through imputation. We developed a validated four-stage framework for imputation specifically designed to preserve biological signal needed for digital twin development, while quantifying trade-offs in downstream analyses. Using retinal imaging and omics data from the NASA RR9 mission as a case study (9), we demonstrate how to diagnose why data is missing (10), select and optimize appropriate imputation strategies (5,10), and rigorously evaluate whether imputed data remains biologically meaningful. A key finding of this work is that while imputation substantially improves the performance of predictive models, it can simultaneously obscure subtle biological patterns; a critical trade-off that researchers must understand before applying these methods (11). This framework provides practical, actionable guidance for space biologists and data scientists working with sparse, multimodal datasets in space biology, and represents a foundational step toward more complete and reliable data-driven models of human physiology in extreme environments.

## 3. Background

### 3.1 Imputation Techniques

Imputation is a fundamental step in data analysis that involves reconstructing missing values by learning from relationships within a dataset, thereby enabling downstream analyses and models to operate on a complete dataset without discarding samples or features (12). In the context of small, high-dimensional, and multimodal datasets characteristic of space biology, the chosen imputation strategy can materially alter downstream significance patterns, correlation structures, network topology, and model calibration. Suboptimal imputation choices may attenuate associations, bias parameter estimates, and underestimate uncertainties, with significant downstream impacts on both inference and prediction.

Imputation methods broadly fall into four categories, which are often used in combination:

1. **Simple methods**: These include basic replacement strategies such as the mean, median, or mode for tabular features. While straightforward and computationally inexpensive, these methods do not account for relationships between variables and can introduce bias in heterogeneous datasets (13).
2. **Statistical models**: This category encompasses regression-based imputation and expectation-maximization algorithms. These methods leverage aggregate patterns in data and are most effective under well-characterized missingness mechanisms (14).
3. **Machine learning methods**: These algorithms learn patterns from observed data to better estimate missing values, capturing more complex relationships than simple statistical models. Key approaches include:

○ **k-Nearest Neighbors (kNN) imputation**, which identifies the k most similar samples to the one with missing data and uses their values to impute the missing entry. kNN has intuitive appeal in leveraging data continuity but can struggle in high-dimensional datasets or when missingness occurs in variables without close neighbors (15,16).

○ **Multiple Imputation by Chained Equations (MICE)**, an iterative approach that fills missing data by modeling each feature with missing values conditional on all other features in the dataset, progressively improving imputations across iterations (17,18).

○ **Random Forests (MissForest)**, a non-parametric iterative imputation algorithm that uses an ensemble of decision trees to predict missing values, capturing non-linear interactions and performing well on mixed data types (19).

○ **Deep learning-based methods**, including denoising autoencoders, generative adversarial imputation networks, and diffusion models, offer state-of-the-art performance for complex biomedical data but require large training datasets and substantial computational resources (20,21).

1. 4. **Domain-specific rules**: These involve utilizing known biological, physical, or operational constraints to handle missing information by design, and are particularly useful when the structure of missingness is well understood (22).

### 3.2 Missingness Mechanisms

The missingness mechanism is a crucial factor that should guide the selection of an appropriate imputation method (23). Three primary mechanisms are recognized:

● **Missing Completely at Random (MCAR)**: Missingness is independent of both observed and unobserved values, such as a lab instrument malfunction causing random omissions. Simple or model-based methods are least biased in this scenario (24,25).
● **Missing At Random (MAR)**: Missingness depends on observed variables but not on the unobserved value itself. This is the most common mechanism in space biology datasets from the NASA Open Science Data Repository, where experimental design choices and tissue allocation constraints systematically drive which measurements are available (26).
● **Missing Not at Random (MNAR)**: Missingness depends on the unobserved value itself, such as astronauts with severe vision changes having incomplete ocular scans precisely because of the severity of their condition. This is the most challenging mechanism to address and can introduce systematic bias if not properly accounted for (25,27).

Since real datasets rarely conform to a single missingness mechanism, careful diagnosis is essential before selecting an imputation strategy.

### 3.3 Rationale for Methods Selected

Given the MAR missingness mechanism identified in the RR9 dataset and the multimodal, heterogeneous nature of the data, we selected imputation methods that leverage inter-variable relationships rather than simple univariate approaches (5).

We implemented three strategies:

1. KNN imputation
2. MICE with ElasticNet regularization, and
3. Per-column hybrid strategy that selects between KNN and MICE independently for each feature based on empirical performance.

These methods were chosen for their ability to model complex dependencies between features, their compatibility with mixed data types, and their scalability to high-dimensional omics data (28). A random sample imputer served as a computationally inexpensive baseline. Deep learning methods were not implemented due to small sample size (N=20 per group), which is insufficient for reliable model training without overfitting risk (29).

For the hybrid strategy, we selected KNN and MICE based on empirical performance: KNN achieved the lowest overall masked value recovery Root Mean Squared Error (RMSE = 11.20) and excelled on morphological and low-dimensional features, while MICE (RMSE=22.80) excelled on high-dimensional omics with complex inter-feature dependencies. The hybrid approach leverages these complementary strengths by selecting the best-performing method per feature. Detailed hyperparameter optimization procedures for both methods are provided in Section 4.2.

## 4. Methods

### 4.1 Data Integration and Exploratory Analysis

#### RR9 Mission Background

The Rodent Research 9 (RR-9) payload consisted of three space biology investigations examining the impacts of long-duration spaceflight on visual impairment, fluid shifts, and joint tissue degradation. A total of 100 male C57BL/6J mice (10 weeks old at launch) were utilized for the study, distributed evenly into five experimental and control cohorts of 20 mice each. The experimental Flight (F) group consisted of 20 mice maintained in microgravity aboard the International Space Station (ISS) for approximately 35 days before returning to Earth alive. This study focuses exclusively on ocular data collected from this payload, spanning two of the three investigations:

- Investigation 1 (PI: Michael Delp), which examined the effects of microgravity on cephalad fluid shifts and increased intracranial pressure (30), and
- Investigation 2 (PI: Xiao Wen Mao), which characterized the impact of spaceflight on retinal vasculature and blood-retinal barrier integrity (30).

To isolate the biological effects of microgravity from environmental and temporal variables, the control dataset was divided into four distinct Earth-based comparison groups of 20 mice each: Ground Control (GC), Vivarium Control (Viv), Cohort Control 1 (CC1), and Cohort Control 2 (CC2). The GC group was housed in identical hardware simulators mirroring the environmental conditions of the ISS, while the Viv group was kept in standard laboratory caging. Crucially, the original GC and Vivarium experiments at Kennedy Space Center had to be completely cancelled and rescheduled due to the facility evacuation and power hazards caused by Hurricane Irma in September 2017. Because the replicated GC and Vivarium experiments could not be completed until May 2018 using a different generation of mice, the CC1 (2017 birth cohort) and CC2 (2018 birth cohort) groups were introduced as necessary temporal controls to statistically validate that age-matched biological and genetic baselines remained uniform across both animal cohorts (30).

Nine assays were performed on retinal and ocular tissue across the RR9 datasets. Lipid peroxidation was evaluated by immunofluorescence staining with an antibody against 4-hydroxynonenal (HNE) (31). Expression of tight junction protein zona occludens-1 (Zo-1) and platelet endothelial cell adhesion molecule- 1 (PECAM- 1) were evaluated by immunofluorescence staining with antibodies against Zo-1 and CD31/PECAM-1, respectively (32). Cone photoreceptors were detected by immunostaining with peanut agglutinin (PNA) (31). Apoptosis was assessed using a terminal deoxynucleotidyl transferase dUTP nick end labeling (TUNEL) assay, which quantifies DNA fragmentation (31,32). Nuclei were counterstained with DAPI, and for TUNEL, Zo-1, and PECAM assays, the endothelium was additionally stained with lectin . Double staining was performed for HNE and PNA on a single section, whereas Zo-1 and PECAM-1 were stained on separate sections (9). Immunohistochemistry (IHC) quantification was performed using assay-specific metrics, including fluorescence intensity for some stains and positive cell counts reported as density for others. Micro-computed tomography (microCT) images were acquired to capture changes in ocular morphology, including alterations in retinal thickness and surrounding structures; the middle slice of the sagittal view of each sample was used for measurement (32). Intraocular pressure was evaluated by tonometry (32).

Multimodal ocular datasets were accessed from the NASA Open Science Data Repository (OSDR) via the OSDR API. The five datasets used in this study are summarized in Table 1.

**Table 1.** RR9 ocular datasets accessed from the NASA OSDR used in this study.

| Identifier | Assay Types |
| --- | --- |
| OSD-557 | HNE (Immunostaining Microscopy), micro-CT, PNA (Immunostaining Microscopy) |
| OSD-568 | TUNEL, Zo-1 (Immunostaining Microscopy), PECAM (Immunostaining Microscopy) |
| OSD-715 | Proteomics (Protein expression profiling) |
| OSD-255 | RNA-Seq (Transcription profiling) |
| OSD-583 | Tonometry |

**Table 2.** Estimators evaluated during the second stage of imputation.

| Model | Hyper parameters |
| --- | --- |
| ARD Default | Default |
| ARD Sparse | [alpha_1,alpha_2, lambda_1, lambda_2]=1e-4 |
| ARD Dense | [alpha_1,alpha_2, lambda_1, lambda_2]=1e-6 |
| ElasticNet Default | [alpha, l1_ratio, max_iter]=[0.1, 0.5, 500] |
| ElasticNet L1 | [alpha, l1_ratio, max_iter]=[0.1, 0.9, 500] |
| ElasticNet L2 | [alpha, l1_ratio, max_iter]=[0.1, 0.1, 500] |
| ElasticNet Weak | [alpha, l1_ratio, max_iter]=[0.01, 0.5, 500] |
| ElasticNet Strong | [alpha, l1_ratio, max_iter]=[1.0, 0.5, 500] |

### 4.2. Data Observations and Missingness Descriptions

Examination of the data availability patterns across the RR9 dataset suggested that missing data predominantly reflects experimental design choices and tissue allocation constraints rather than random sample loss or experimental failure. The limited size of ocular tissue imposed practical constraints on assay allocation: tissue must be sectioned prior to staining, and unless double stained, each staining protocol requires distinct sections. For protein expression profiling and RNA-seq, tissue must be lysed, and for microCT, tissue must be fixed, dehydrated, and stained. The structured missingness patterns observed across samples are consistent with practical constraints imposed by the limited size of ocular tissue, which would preclude performing every assay on every eye. As a result, samples were systematically distributed across assays such that some mice contributed to specific measurements while others were reserved for different analyses. Assay allocation varied not only across mice but also across eye laterality, such that different assays could originate from the right versus left eye of the same mouse (31,32).

Although a larger number of samples were processed and imaged, for some assays only a subset that would potentially be sufficient for statistical analysis was quantified. For example, in the case of HNE and PNA assays, double staining was performed, yet examination of the data revealed that images from selected samples in certain control groups were not analyzed for PNA (32). Consequently, some missing values correspond to unquantified measurements rather than unassayed or lost samples.

An exception is tonometry, which was performed as an in vivo measurement and therefore did not require tissue harvesting or sectioning. Tonometry data were therefore acquired for all available samples, with no analogous constraints on tissue availability (31).

### Nullity Correlation Analysis and Hierarchical Clustering of Missingness

Multi-modal RR9 retinal datasets spanning nine assay types across tonometry, RNA-seq, proteomics, Zo-1 immunostaining, TUNEL, PECAM, microCT, PNA, and HNE, were harmonized on a shared sample identifier (Source Name) and merged by outer join into a single sample-by-feature matrix. Features within each assay were grouped into assay-level categories prior to missingness analysis.

Missingness was evaluated at the assay-category level rather than at the individual feature level. For each sample and each assay category, a binary missingness indicator was defined using a coarse aggregation rule: a sample was classified as missing for a given assay if any feature within that assay category was null (NaN). Conversely, a sample was classified as present for an assay if at least one feature in that category was observed. This .any() rule treats partial and complete absence equivalently - a sample with one missing protein measurement is flagged the same as a sample missing the entire proteomics panel. Sensitivity to the .any() threshold was not assessed; alternative rules (e.g., requiring all features to be missing) would yield stricter missingness definitions.The approach prioritizes sample-level assay availability over feature-level completeness and is appropriate for identifying whether entire assay suites were collected for a given animal, but it does not distinguish degrees of within-assay missingness.

From these binary indicators, two complementary visualizations were generated. First, a nullity correlation matrix (38) was computed as the Pearson correlation between binary missingness indicators across samples for each pair of assay categories. Positive correlations indicate co-missingness (samples missing one assay tend to miss the other); values near zero indicate independent missingness; negative values indicate weak inverse association. The correlation heatmap was generated using the Python missingno library (39), with tonometry excluded from correlation analysis because it exhibited no missing values (zero variance in the missingness indicator). Assay categories with no missing values produce undefined correlations and were assigned a correlation of zero for display purposes in the full matrix. Second, To assess hierarchical structure in missingness patterns, a nullity dendrogram was generated using msno.dendrogram, which applies average-linkage hierarchical clustering to binary nullity profiles across samples. Assays that merge at lower linkage distance share more similar missingness patterns across animals. Both visualizations used the same category-level binary encoding described above (38,39).

This category-level framework was chosen because RR9 assays differ substantially in feature dimensionality (e.g., proteomics comprises thousands of features versus a handful of imaging readouts) and because missingness in this dataset primarily reflects sample-level assay dropout rather than sporadic feature-level gaps.

### Missingness Mechanism Diagnosis

To diagnose the underlying missingness mechanism, a logistic regression model was trained for each data category to predict its missingness indicator vector (1=missing, 0=present) using the missingness indicators of all other categories as features. This approach was inspired by the principles of Little’s MCAR test (40), evaluating whether missingness in any given category was predictable from the observed data structure of other categories.

### Exploratory Principal Component Analysis

To assess the inherent structure and separability of experimental groups, PCA (33) was applied individually to several data modalities, including the full RNA-seq dataset, a curated subset of 84 phenotype-related genes, proteomics, microCT, TUNEL assay data, and HNE immunostaining data. An additional PCA was performed on a combined dataset of four immunostaining assays (Zo-1, PECAM, PNA, and HNE).

### Phenotypically Relevant Genes Identification

Due to the large feature dimensions of the RNA-seq data (n=23,456) and the redundant nature of gene expression data, we identified a subset of genes with high predictive power for the phenotype measurements. Phenotype-related gene sets were first collected from Harmonizome and MSigDB (34,35), providing a large list of genes nominally related to each phenotype. Human gene IDs from Harmonizome were converted to mouse IDs. For each phenotype, the data were subset to include only samples with measurements from both RNA-seq and the target phenotype. The top 100 genes most correlated with each phenotype were selected, and an XGBoost regression model was trained using cross-validation to identify genes with high predictive power (36). This process resulted in a final subset of 26 genes with high predictive power for the phenotypic outcomes measured in this study.

### Correlation Analysis for Biological Anchoring

To anchor the imputation of RNA-seq and proteomics data to biological characteristics, Pearson correlations were calculated between RNA-seq data and both HNE and TUNEL assay data for the subset of samples where these measurements overlapped (37).

### 4.3. Data Imputation Strategies

Imputation was applied to recover measurements for all 100 samples in the RR9 experimental design, of which only a subset had measurements available for any given assay due to tissue allocation constraints. Based on the missingness mechanism identified as MAR, several multivariate imputation techniques were implemented (5,18). As a critical preparatory step for all methods, the dataset’s feature columns were optimally reordered based on correlation-based hierarchical clustering to enhance the stability and performance of the iterative imputation algorithms (5,41). This process arranges columns with similar missingness patterns adjacently, ensuring that the most relevant information is available when predicting a missing value. The methodology was as follows:

1. A binary nullity matrix was created for all numeric feature columns, where an entry was set to 1 if the data point was missing and 0 otherwise. Identifier columns (Source Name, Group) were excluded from this process.
2. The pairwise Pearson correlation of this nullity matrix was computed and transformed into a distance matrix using the formula distance = 1 − |correlation|, ensuring that features with highly correlated missingness patterns have a small distance between them.
3. Agglomerative hierarchical clustering with average linkage was performed on the condensed distance matrix and visualized as a truncated dendrogram showing the top 25 clusters for clarity.
4. The optimal leaf ordering was extracted from the resulting linkage matrix. The dataset used for imputation was then reconstructed with identifier columns placed first, followed by the optimally ordered feature columns.

This structured ordering places features with the most similar missingness patterns next to each other, a crucial step designed to optimize the performance of all subsequent iterative imputation algorithms (42).

### Group Encoding

Prior to imputation, the categorical Group variable was encoded as an ordinal variable reflecting the biological proximity of each control subgroup to the spaceflight condition (43). Flight samples were assigned a value of 0, ground control samples (GC) a value of 1, cohort control group 1 (CC1) a value of 2, cohort control group 2 (CC2) a value of 3, and vivarium control samples (Viv) a value of 4. This encoding preserves the biological ordering of experimental groups from spaceflight conditions through increasingly standard laboratory conditions (44). This provides the imputation algorithm with a continuous representation of group membership that is more informative than binary or one-hot encoding. Binary encoding (Flight=1, all others=0) was evaluated as an alternative but yielded substantially higher imputation RMSE (76.11 vs 11.20), confirming that ordinal encoding better captures the structured relationship between experimental groups for the purposes of imputation.

### K-Nearest Neighbors (KNN) Imputer

The KNN imputer was optimized using a grid search approach based on masked value recovery on the full multimodal dataset. The ordinal-encoded Group variable was appended as an additional numeric feature column prior to imputation, providing group membership information to the imputer in a continuous form that reflects the biological ordering of experimental conditions.

To evaluate hyperparameter combinations, 10% of observed non-missing values across the full dataset were randomly masked to serve as a held-out test set. The dataset was then standardized using column-wise mean and standard deviation computed from the masked dataset, with zero-variance columns assigned a standard deviation of 1 to prevent division by zero. A grid search was performed across combinations of n_neighbors (2, 3, 5, 7, 10) and weights (uniform, distance), evaluating each combination by computing the RMSE between true and imputed values at the masked positions after inverse-transforming the imputed output back to the original scale.

The optimal parameters were identified as n_neighbors=5 and weights=uniform, yielding a masked recovery RMSE of 11.20. These parameters were then applied to impute the full multimodal dataset. The fitted scaling parameters (column means and standard deviations) and imputation pipeline were saved for reproducibility and subsequent validation analyses.

### Random Sample Imputer

A random sample imputer was applied as a computationally inexpensive baseline method (45). This approach replaces missing values by randomly sampling from the observed values of each feature. The process involved a pipeline that scaled the data, applied the imputer, and inverse-transformed the results to the original scale.

### Multiple Imputation by Chained Equations (MICE)

MICE was implemented using the IterativeImputer from scikit-learn with a systematically optimized estimator and hyperparameter configuration (18),(46). Prior to imputation, the Group variable was ordinal-encoded using the same scheme described in Section 4.2 (F=0, GC=1, CC1=2, CC2=3, Viv=4) and appended as an additional numeric feature column.

To identify the optimal MICE configuration for this dataset, a first-stage grid search was performed across 60 parameter combinations: three regression estimators (HuberRegressor, ElasticNet, and ARDRegression), five n_nearest_features values (2, 3, 5, 7, 10), and four maximum iteration counts (3, 5, 7, 10), with initial strategy fixed at mean. For each combination, 10% of observed non-missing values across the full dataset were randomly masked and used as a held-out test set. Each configuration was evaluated by computing the Root Mean Squared Error (RMSE) between true and imputed values at the masked positions after inverse-transforming back to the original scale.

The three estimators were selected to represent a range of regression approaches suited to the biological characteristics of the data:

● **HuberRegressor** — a robust estimator with reduced sensitivity to outliers, appropriate for biological data with heavy-tailed distributions (47).
● **ElasticNet** — a combined L1/L2 regularized estimator providing automatic feature selection, useful in high-dimensional omics settings (48).
● **ARDRegressor** — an Automatic Relevance Determination model that performs sparse Bayesian regression, adaptively down-weighting irrelevant features (49).

Based on first-stage findings, HuberRegressor was excluded from the second-stage search due to systematic lbfgs convergence failures at n_nearest ≥ 7, which produced highly variable and generally poor RMSE values. A focused second-stage grid search was performed on ElasticNet and ARD estimators, evaluating three ARD prior configurations (default, [alpha_1,alpha_2, lambda_1, lambda_2]=1e-4, [alpha_1,alpha_2, lambda_1, lambda_2]=1e-6) and five ElasticNet regularization configurations ([alpha, l1_ratio, max_iter]=[0.1, 0.5, 500], [0.1, 0.9, 500], [0.1, 0.1, 500], [0.01, 0.5, 500], [1.0, .5, 500]) across n_nearest values of 5 and 7 and max_iter values of 3, 5, 10, and 15.

**Table 7.** Top 10 configuration from second-stage MICE grid search.

| Estimator | n_nearest | max_iter | RMSE |
| --- | --- | --- | --- |
| ElasticNet(weak, $\alpha=0.01$) | 5 | 10 | **22.80** |
| ElasticNet (L2, $\alpha=0.1$) | 5 | 10 | 54.35 |
| ElasticNet (default, $\alpha=0.1$) | 5 | 10 | 61.50 |
| ElasticNet (weak, $\alpha=0.01$) | 5 | 15 | 64.03 |
| ElasticNet (L2, $\alpha=0.1$) | 5 | 15 | 64.31 |
| ElasticNet (L1, $\alpha=0.1$) | 5 | 10 | 66.94 |
| ElasticNet (default, $\alpha=0.1$) | 5 | 15 | 68.38 |
| ElasticNet (L1, $\alpha=0.1$) | 5 | 15 | 69.20 |
| ARD (sparse prior) | 5 | 15 | 69.68 |
| ARD (dense prior) | 5 | 15 | 71.51 |

The configuration achieving the lowest masked recovery RMSE was selected as one of the final imputation models (50). The IterativeImputer was configured with the following settings:

● imputation_order=’roman’ — left-to-right column ordering, aligned with the optimal column ordering based on hierarchical clustering.
● skip_complete=True — bypasses columns with no missing values for computational efficiency
● tol=0.01 — convergence tolerance
● initial_strategy=’mean’ — initial fill strategy before iterative refinement
● random_state=42 — for reproducibility

The fitted scaling parameters (column means and standard deviations computed from the full dataset) and the complete imputation pipeline were saved as a joblib file for reproducibility and subsequent validation analyses.

### Cascading Hybrid Imputation Strategy

A per-column hybrid imputation strategy was developed to maximize imputation accuracy by selecting the best-performing method for each feature independently, rather than applying a single method uniformly across the dataset. This approach leverages the complementary strengths of KNN and MICE with weak ElasticNet regularization, which were identified as the two best-performing methods from the individual grid searches.

Two validated imputation pipelines were used as candidate methods:

● **KNN** (n_neighbors=5, weights=uniform, ordinal group encoding) — optimal for high-dimensional features with local neighborhood structure
● **MICE with weak ElasticNet** (alpha=0.01, l1_ratio=0.5, n_nearest=5, max_iter=10, ordinal group encoding) — optimal for features with linear inter-variable relationships.

To enable honest per-column method comparison, both pipelines were re-run on a masked version of the original dataset with 10% of observed values held out (random seed=42, identical to the individual method evaluations). Per-column RMSE was computed in z-score space for each method at the held-out positions. For each column with missing values, the method achieving the lower RMSE on the held-out values was selected to fill the missing entries from the corresponding pre-computed full-data imputation. For columns with no held-out values, MICE with weak ElasticNet was used as the fallback method.

The final hybrid imputed dataset was assembled by combining the selected imputed values for each column from the appropriate pipeline. Method selection assignments and per-column RMSE values were saved alongside the final dataset for full reproducibility and interpretability.

### 4.4. Imputation Validation Framework

A rigorous, multi-faceted framework was established to evaluate the quality and analytical utility of each imputed dataset compared to the original unimputed data. All validation steps were performed on cleaned datasets subsetted to intersecting numeric RNA-seq features.

### Supervised Classification

The impact of imputation on downstream predictive modeling was assessed by training Support Vector Machine (SVM) and Random Forest (RF) classifiers to distinguish between flight and non-flight samples. Performance was evaluated using five-fold cross-validation accuracy and held-out test set accuracy for both classifiers. Receiver Operating Characteristic (ROC) curves were additionally generated for the SVM classifier to assess discriminative ability across imputation methods(51),(52).

### Unsupervised Clustering

K-means clustering (k=2) was performed to assess the preservation of latent group structures in each imputed dataset. The agreement between the resulting clusters and the true binary flight/ non-flight labels was quantified using the Adjusted Rand Index (ARI), where a value of 1 indicates perfect agreement and a value of 0 indicates agreement no better than chance (52). PCA was used to project the original and imputed datasets into two dimensions to visually compare the separation of flight and non-flight clusters across imputation methods.

### Correlation Structure Preservation

To assess whether imputation preserved the inter-feature correlation structure of the original data, the Spearman correlation matrix was computed for both the original and each imputed dataset (52). The Spearman correlation coefficient between the two matrices was then calculated and reported as a summary measure of correlation structure fidelity.

### Marginal Distribution Similarity

The Kolmogorov-Smirnov (KS) statistic was calculated between the original and imputed distributions for each feature to measure distributional fidelity (37). The average KS statistic across all features was reported, where a value of 0 indicates identical distributions and a value of 1 indicates completely non-overlapping distributions.

### Masked Value Recovery

To directly measure imputation accuracy independently of sample size and classifier choice, a masked value recovery analysis was performed on the original 16-sample RNA-seq dataset. 20% of the observed values were randomly masked to simulate missing data. Each imputation method was then applied to reconstruct the masked values, and the imputed values were compared against the true value using RMSE and Pearson correlation coefficient. This approach provides a direct, ground-truth evaluation of imputation accuracy that is not confounded by the sample size disparity between the origins and imputed datasets (50).

### Cross-Dataset Validation

To assess whether imputer datasets produce classifiers that generalize to real observed data, each imputed dataset was used to train SVM and Random Forest Classifiers, which are then tested on the original 16 observed samples as a completely held-out test set. This train-on-imputed, test-on-original design directly evaluates whether imputation-derived models capture genuine biological signal rather than artifacts introduced by the imputation process (23). Five-fold stratified cross-validation was additionally performed on the training set to assess classifier stability.

### Permutation Test

To validate that classifier results were attributable to chance, a permutation test was performed on the SVM classifier for each dataset (5). Flight/non-flight labels were randomly shuffled 1000 times, and the classifier was re-trained and evaluated on each permutation to generate a null distribution of accuracy scores. The true classifier accuracy was compared against this null distribution, and a p-value was computed as the proportion of permuted scores exceeding the true score. A significant result (p<0.05) confirms that the classifier is learning a genuine biological signal rather than noise.

### Visualization

PCA was used to project the original and imputed datasets into two dimensions to visually compare the separation of flight and non-flight clusters. ROC curves were generated for the SVM classifiers to assess discriminative ability across imputation methods. Scatter plots of true versus imputed values were generated for the masked value recovery analysis. Histograms of permuted accuracy scores with the true score marked were generated for the permutation test.

## 5. Results

### 5.1 Data Observations and Missingness Descriptions

#### Dataset Overview

Initial exploration of the RR9 dataset revealed substantial heterogeneity in sample size and feature dimensionality across the nine assay categories. Tonometry was the most densely sampled modality with 100 samples, while immunostaining assays ranged from 10 to 23 samples. RNA-seq and proteomics datasets contained 16 and 12 samples respectively, with high feature dimensionality (23,419 and 6,634 features). The full sample and feature counts for each modality are summarized in Table 3. The data availability heatmap (Figure 2) reveals the structured nature of missingness across experimental groups, with tonometry being the only modality present across all samples and groups, while immunostaining assays show group-specific patterns of availability.

**Figure 1.**
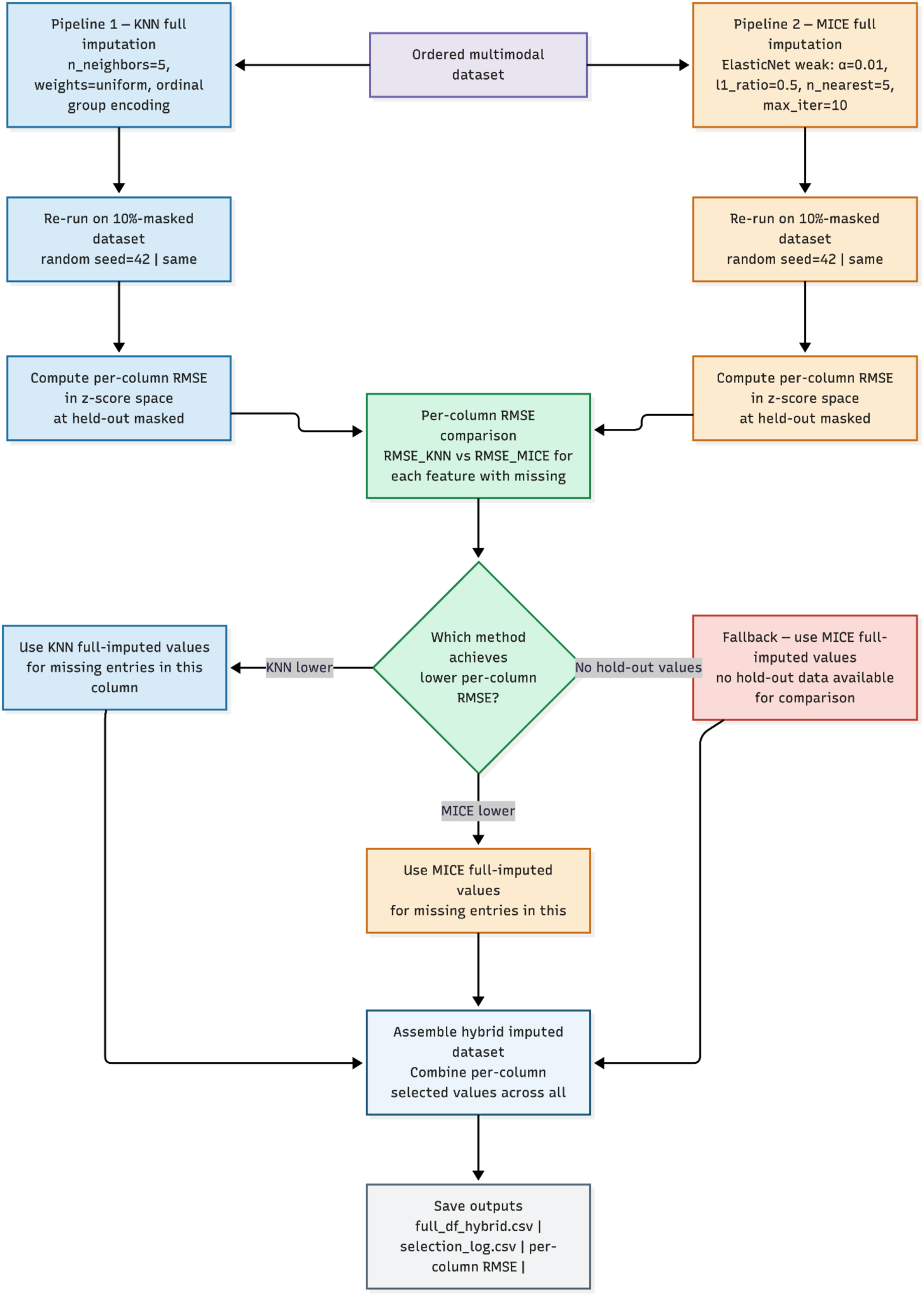
Per-column Hybrid Imputation Strategy. Two candidate imputation pipelines - KNN (n_neighbors=5, weights=uniform) and MICE with ElasticNet regularization (alpha=0.01, l1_ratio=0.5, n_nearest=5, max_iter=10) - are both applied to the full dataset and independently re-run on a version of the data with 10% of observed values randomly masked. Per-column RMSE is computed in z-score space for each method at the masked positions. For each feature with missing values, the method achieving the lower masked value recovery RMSE is selected to fill the missing entries from the corresponding pre-computed full-data imputation. For columns with no held-out values available for comparison, MICE with ElasticNet is applied as the fallback method. The final hybrid imputed dataset is assembled by combining the per-column selected values and saved alongside a full method selection log and per-column RMSE summary for reproducibility.

**Figure 2.**
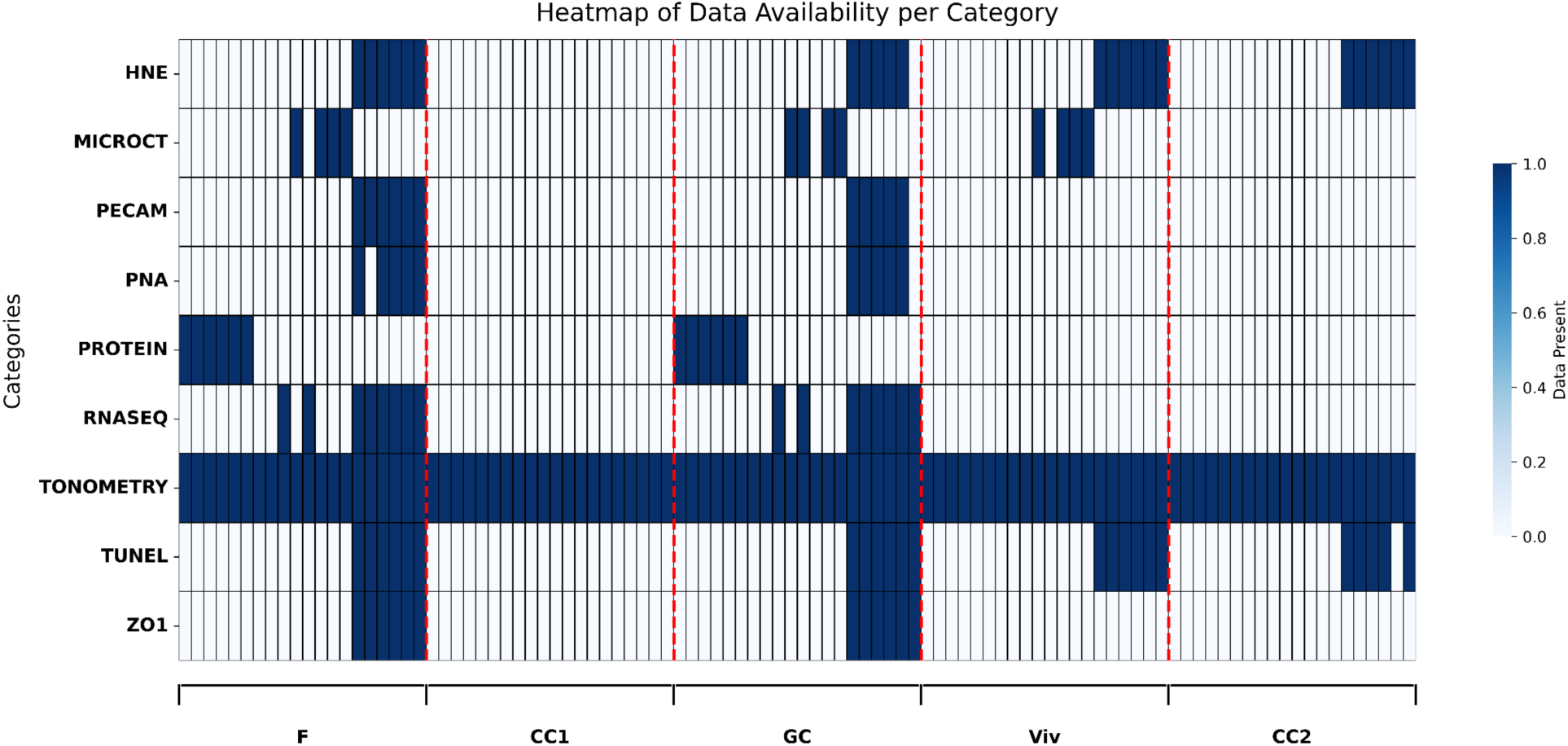
Heatmap of Data Availability per Category. Each row represents a data category and each column represents an individual sample, grouped by experimental condition (F, CC1, GC, Viv, CC2). Dark blue indicates data present; white indicates missing data. Red dashed lines separate experimental groups. Tonometry is the only category with complete data across all samples and groups, while immunostaining assays exhibit structured, group-specific missingness patterns.

**Table 3.** Sample size and feature dimensionality across RR9 ocular assay categories. For each data modality, the number of samples and measured features are reported. Tonometry is the most densely sampled modality, while immunostaining assays have substantially smaller sample sizes reflecting tissue allocation constraints. RNA-seq and proteomics datasets are characterized by high feature dimensionality relative to sample size, a challenge that motivates the imputation framework described in this study.

| Data | Number of Samples | Number of Features |
| --- | --- | --- |
| Tonometry | 100 | 8 |
| RNA-Seq | 16 | 23,419 |
| Proteomics | 12 | 6,634 |
| Zo-1 | 12 | 1 |
| TUNEL | 23 | 5 |
| PECAM | 11 | 1 |
| microCT | 12 | 12 |
| PNA | 10 | 4 |
| HNE | 23 | 5 |

**Table 4.** Assay pairs with the strongest positive nullity correlations in the RR9 retinal dataset. Values represent Pearson correlation coefficients between binary missingness indicators defined at the assay-category level (missing = any null feature within the assay). Higher values indicate that the same samples tend to lack both assays.

| Assay pair | r |
| --- | --- |
| PECAM ↔ PNA | 0.95 |
| PECAM ↔ Zo-1 | 0.95 |
| HNE ↔ TUNEL | 0.94 |
| PNA ↔ Zo-1 | 0.9 |
| RNA-seq ↔ Zo-1 | 0.85 |
| PECAM ↔ RNA-seq | 0.81 |
| PNA ↔ RNA-seq | 0.76 |

#### Nullity Correlation Analysis and Hierarchical Clustering of Missingness

The nullity correlation heatmap quantified pairwise relationships among assay missingness patterns (Figure 3). Tonometry was absent from the missingness correlation plot because no samples lacked tonometry data.

**Figure 3.**
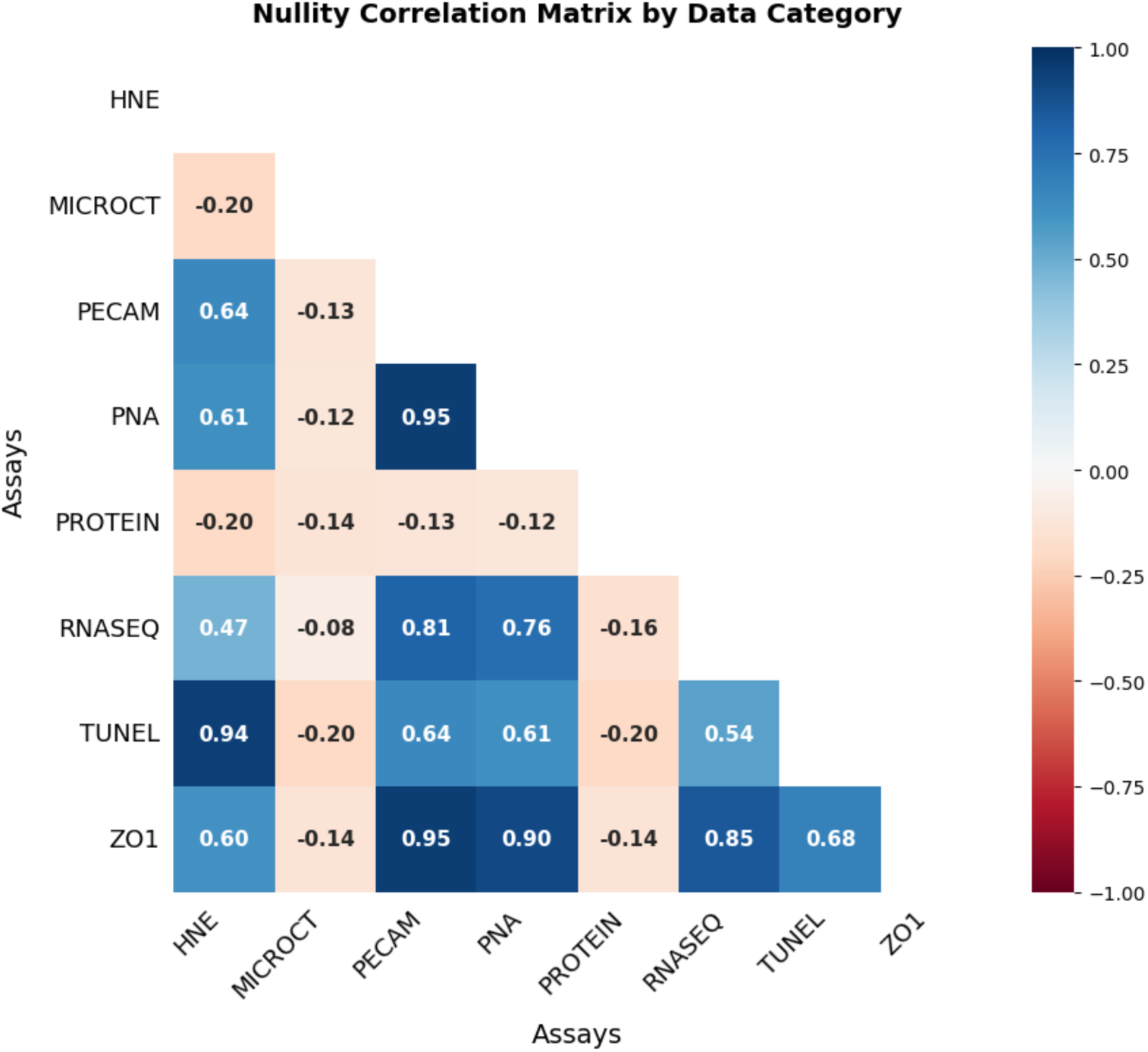
Nullity Correlation Matrix by Data Category. Pearson correlation coefficients between the missingness indicator vectors of each data category pair. Strong positive correlations (dark blue) indicate categories that are systematically absent in the same samples, while near-zero or negative correlations (white to red) indicate independent missingness patterns. Tonometry shows no correlation with any other category, reflecting complete data availability. RNA-seq, ZO1, PECAM, PNA, HNE, and TUNEL form a positive correlated missingness cluster.

Several assay pairs exhibited strong positive co-missingness (Pearson r ≥ 0.75), indicating that the same samples tended to lack multiple assays simultaneously:

These values support a shared sample-level dropout mechanism among PECAM, PNA, Zo-1 RNA-seq, and two other immunohistochemistry (HNE, TUNEL) assays - when one assay in this group was absent for a sample, companion assays were typically absent as well.

Moderate positive correlations (r ≈ 0.47–0.68) linked HNE and TUNEL to the immunostaining/ RNA-seq cluster, suggesting a broader retinal assay block with a tighter HNE–TUNEL sub-structure.

MicroCT and proteomics showed weak near-zero to slightly negative correlations with all other assays (r ≈ −0.08 to −0.20), indicating that their missingness patterns were largely independent of the co-missing retinal assay group. These weak negative values are not large enough to support a meaningful inverse missingness relationship and are best interpreted as absence of shared structure rather than systematic mutual exclusivity.

Together, the two heatmaps (Figure 2 and Figure 3) demonstrate that RR9 missing data are highly structured: completeness varies by cohort, and most assays co-miss at the sample level, while microCT and proteomics follow distinct missingness profiles.

Hierarchical clustering of assay-level missingness profiles (Figure 4) corroborated the correlation heatmap (Figure 3) and resolved assay groupings into a clear tree structure.

**Figure 4.**
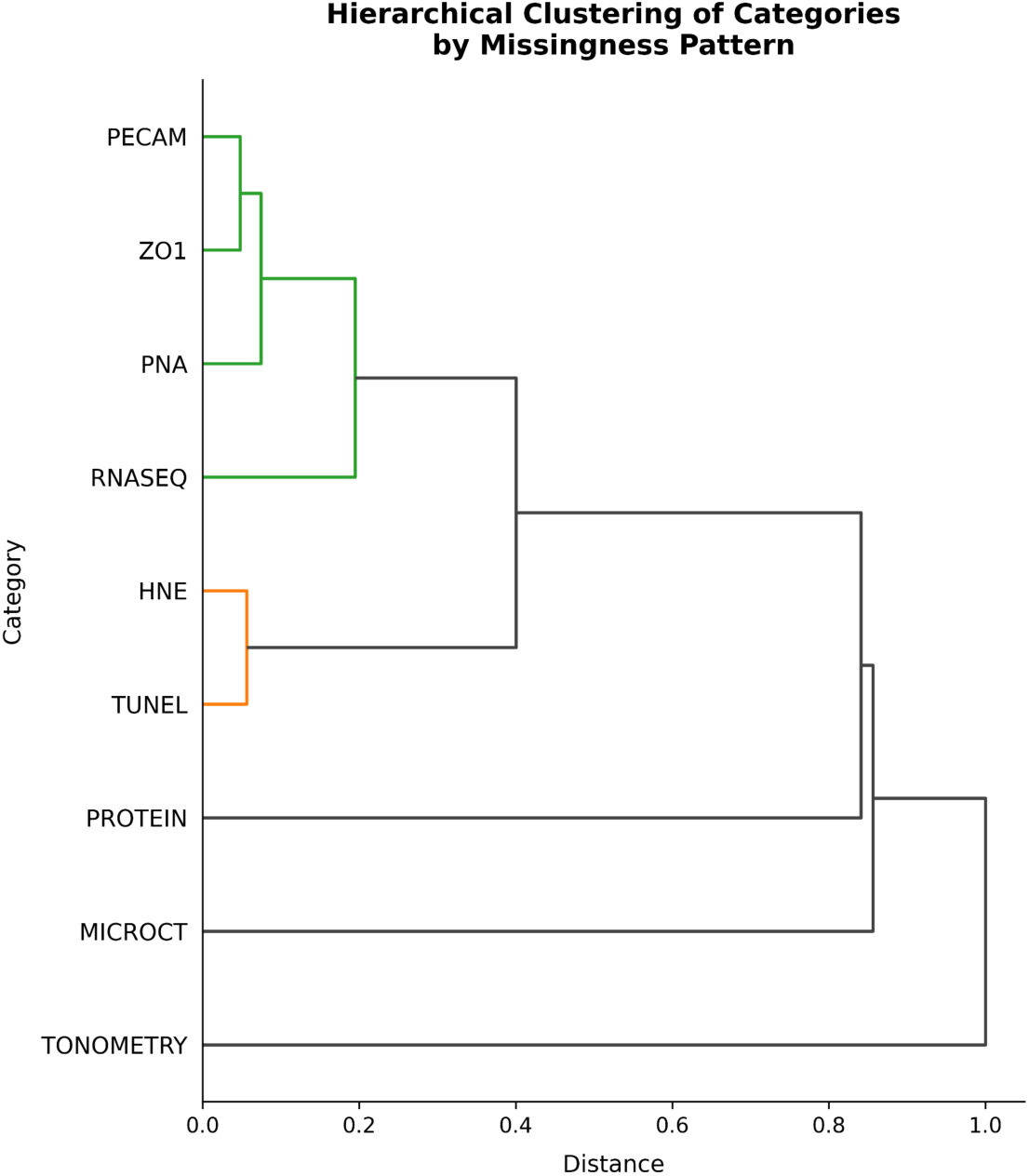
Hierarchical Clustering of Categories by Missingness Pattern. Agglomerative hierarchical clustering dendrogram based on a distance matrix derived from the nullity correlation matrix (distance = 1 − | correlation|). Categories with similar missingness patterns cluster at short distances. Three broad clusters are visible: a tight immunostaining and sequencing cluster (PECAM, ZO1, PNA, RNA-seq), a paired cluster of lipid peroxidation and apoptosis assays (HNE, TUNEL), and independently missing categories (Protein, microCT). Tonometry is fully isolated at maximum distance, reflecting complete data availability.

At the finest level, PECAM and Zo-1 were the most similar pair (lowest merge distance), followed closely by PNA, which joined the PECAM-Zo-1 cluster. RNA-seq subsequently merged with this immunostaining group, forming a core cluster of four assays whose missingness patterns were nearly interchangeable across samples.

HNE and TUNEL formed a separate sub-cluster before joining the larger immunostaining/RNA-seq group at a higher linkage distance. This hierarchy is consistent with a two-tier structure: HNE and TUNEL co-miss most strongly with each other, while still sharing broader missingness patterns with the PECAM/PNA/Zo-1/RNA-seq block.

MicroCT and proteomics were the most dissimilar assays. They clustered with each other only at the highest linkage distances and joined the full tree last, confirming that their missingness patterns were weakly related both to each other and to the retinal assay super-cluster.

In summary, the dendrogram supports the conclusion that the same animals tend to be missing similar retinal assay data, particularly among PECAM, PNA, Zo-1, RNA-seq, HNE, and TUNEL, whereas microCT and proteomics missingness appear largely assay-specific. This structured missingness has direct implications for downstream analysis: linked assay groups should be evaluated and imputed jointly, while microCT and proteomics warrant separate treatment.

#### Missingness Mechanism Diagnosis

Logistic regression models trained to predict missingness indicator of each category from the missingness indicators of all other categories achieved high predictive accuracy in all categories (40), ranging from 0.88 (microCT, Protein) to 1.0 PECAM, as summarized in Table 5. This high predictability provides evidence against Missing Completely At Random (MCAR) mechanism and supports Missing At Random (MAR) mechanism, in which the probability of missingness in any given category is highly dependent on the observed missingness patterns of other categories. Tonometry was excluded from this analysis as it contained no missing values. These findings confirm that multivariate imputation methods capable of leveraging inter-variable relationships are appropriate for this dataset, and that simple univariate approaches such as mean or median imputation are not recommended.

**Table 5.** Logistic Regression-Based Missingness Mechanism Diagnosis for Each Data Category. For each category with missing data, a logistic regression model was trained to predict its missingness indicator from the missingness indicators of all other categories. Model accuracy reflects the degree to which missingness in a given category is predictable from observed data structure in other categories. High accuracy values provide evidence for a Missing At Random (MAR) mechanism, justifying the use of multivariate imputation methods.

| Category | Skipped | Model Accuracy | Missingness Mechanism Suggestion | Imputation Recommendation |
| --- | --- | --- | --- | --- |
| Tonometry | Yes | - | Complete data (no missingness) | None needed |
| HNE | No | 0.98 | MAR | Use multivariate methods (e.g., MICE) |
| microCT | No | 0.88 | MAR | Use multivariate methods |
| PECAM | No | 1.00 | MAR (perfectly explained) | Use multivariate methods |
| PNA | No | 0.99 | MAR | Use multivariate methods |
| Protein / Proteomics | No | 0.88 | MAR | Use multivariate methods |
| RNA-Seq | No | 0.95 | MAR | Use multivariate methods |
| TUNEL | No | 0.98 | MAR | Use multivariate methods |
| Zo-1 | No | 0.99 | MAR | Use multivariate methods |

#### Exploratory Principal Component Analysis

PCA was performed across all data modalities to assess the inherent structure and group separability of the dataset prior to imputation. The TUNEL assay showed the strongest separation between flight and non-flight groups (Figure 5), with PC1 capturing 73.6% of total variance and 3 components required to exceed the 95% cumulative variance threshold. Flight samples were clearly displaced along PC1 relative to all control groups. The full RNA-seq dataset required 14 components to reach the 95% threshold, with PC1 and PC2 capturing only 10.1% and 9.5% of variance respectively, reflecting the high dimensionality and redundancy of the transcriptomic data. PCA of the predictive gene subset (n=26) substantially improved the structure, with PC1 capturing 30.4% of variance and meaningful separation between flight and ground control samples visible across the first three components (Figure 6).

**Figure 5.**
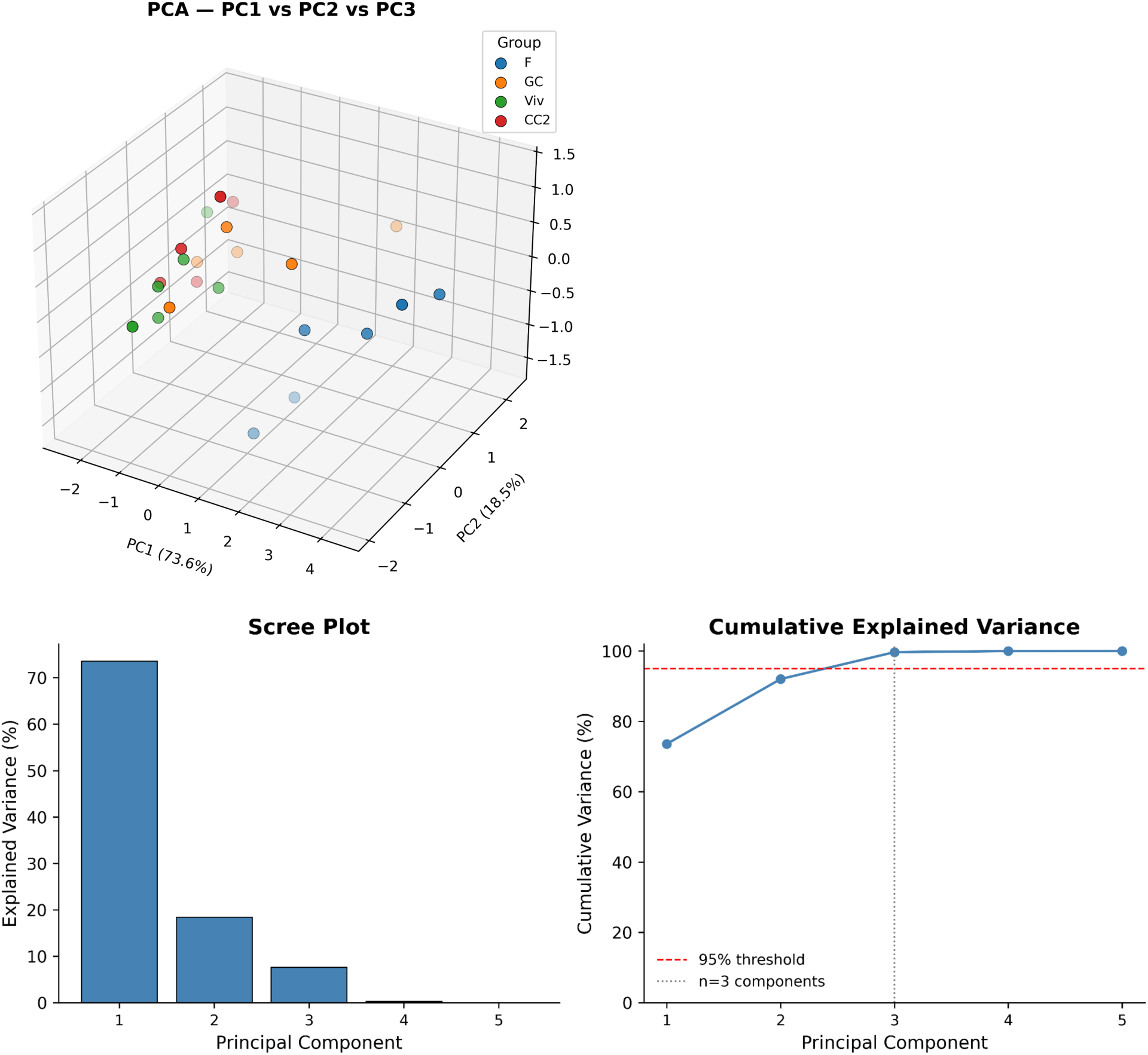
PCA of TUNEL Assay Data. Three-dimensional scatter plot of PC1, PC2, and PC3 colored by experimental group (F, GC, Viv, CC2). PC1 captures 73.6% of total variance. Flight samples are clearly separated from control groups along PC1, indicating that TUNEL-based DNA fragmentation measurements provide the strongest group discriminability among all assay modalities examined. The scree plot and cumulative explained variance plot confirm that 3 components are sufficient to capture 95% of total variance.

**Figure 6.**
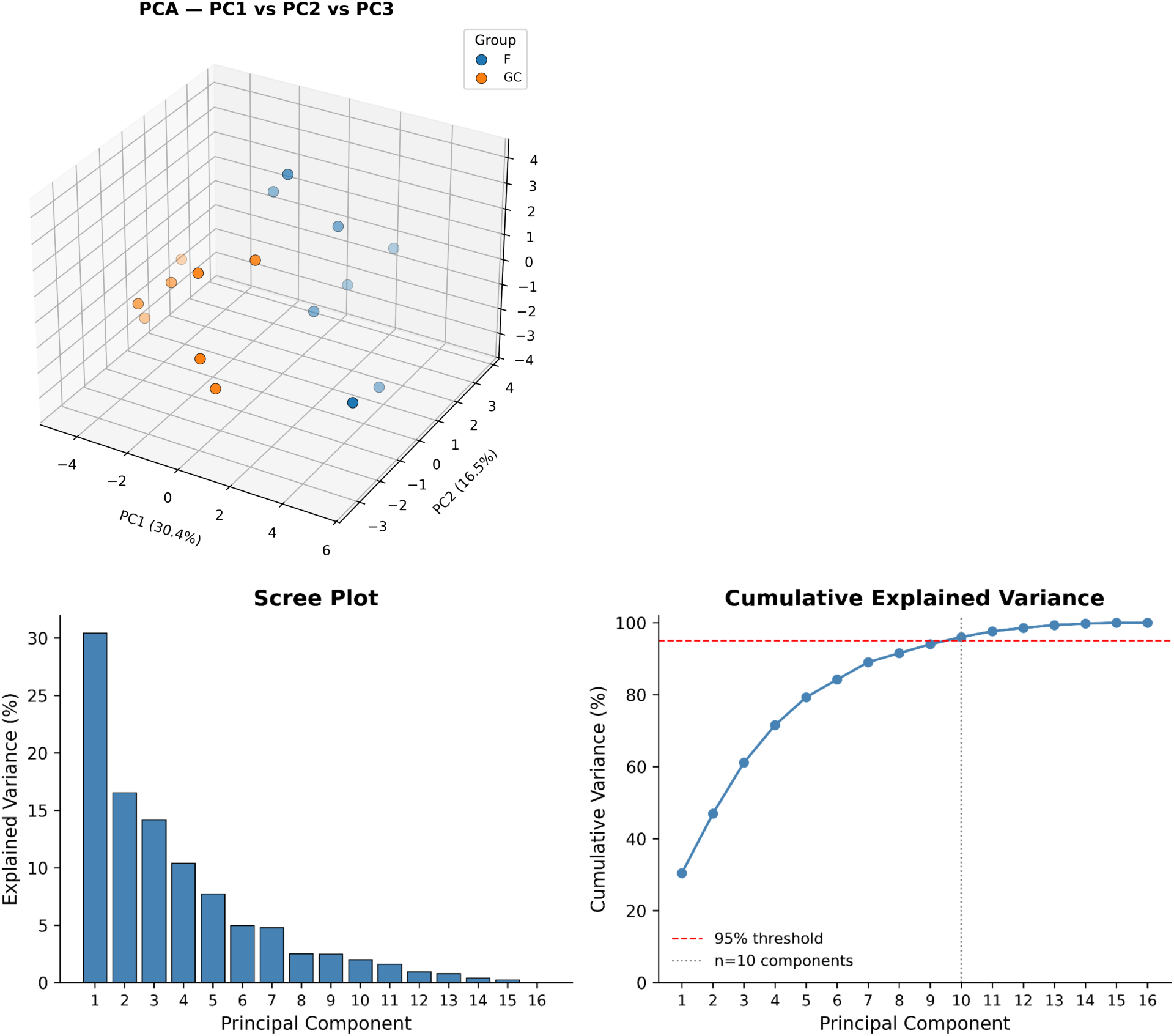
PCA of RNA-seq Predictive Gene Subset. Three-dimensional scatter plot of PC1, PC2, and PC3 for the 26-gene predictive subset colored by experimental group (F, GC). PC1 captures 30.4% of variance, with flight and ground control samples showing partial separation across the first three components. The scree plot confirms that 10 components are required to reach the 95% cumulative variance threshold, reflecting the moderate dimensionality of this curated gene subset.

#### Results of Identifying Gene Highly Predictive of Key Phenotypes

We identified a subset of 26 genes with the highest predictive power across 9 phenotypic measurements, starting from phenotype-related gene sets sourced from Harmonizome and MSigDB, then downselecting via Pearson correlation and XGBoost regression (Table 6). PNA had the strongest predictive relationship with gene expression (n=2 genes; R²=0.86), followed by tonometry measurements (n=9 genes; R²=0.78). Several phenotypes including TUNEL and HNE showed moderate predictive success (R²=0.72 and R²=0.68 respectively), while Zo-1 measurements showed weaker performance (R²=0.38). Notably, right eye tonometry and PECAM measurements showed negative R² values, indicating that the gene sets related to these phenotypes have low overall predictive power. The absence of a predictive gene signal for right eye tonometry is consistent with the original analysis of these data, which reported no significant differences in right eye intraocular pressure measurements following spaceflight. Three phenotypes - retina, choroid, and pigment layer microCT measurements - had insufficient overlapping samples for analysis, with only one sample each having both RNA-seq and phenotype measurements available.

**Table 6.** Results of XGBoost regression analysis to identify predictive genes for each phenotype. R² values reflect the predictive power of the selected gene subset for each phenotypic outcome. Negative R² values indicate that the model performs worse than a horizontal mean baseline, suggesting low gene-phenotype predictive relationships for those measurements.

| Phenotype | R2 from XGBoost | Predictive Genes | N samples with both RNA-seq and Phenotype | Status |
| --- | --- | --- | --- | --- |
| Average_zo1 | 0.38 | 3 | 12 | Analyzed |
| Average_pecam | -6.06 | 2 | 11 | Analyzed |
| Density_EC_tunel | 0.72 | 3 | 12 | Analyzed |
| Retina_microct |  |  | 1 | Skipped - Insufficient samples |
| Choroid_microct |  |  | 1 | Skipped - Insufficient samples |
| Pigment Layer_microct |  |  | 1 | Skipped - Insufficient samples |
| density_pna | 0.86 | 2 | 10 | Analyzed |
| sumcount_hne | 0.68 | 3 | 11 | Analyzed |
| Avg_Left_tonometry | 0.78 | 9 | 16 | Analyzed |
| Avg_Right_tonometry | -3.57 | 4 | 16 | Analyzed |

#### Correlation Analysis for Biological Anchoring

To assess the biological relevance of the identified predictive gene subset, Pearson correlations were computed between RNA-seq expression data and HNE immunostaining measurements for overlapping samples in the Flight and Ground Control groups. Correlation matrices computed using the final 26 predictive genes identified via XGBoost regression revealed strong and structured inter-feature relationships in both the Flight (Figure 7) and Ground Control (Figure 8) groups, supporting the biological relevance of this curated gene subset and HNE assay for anchoring downstream imputation. Notably, the correlation structure differed between Flight and Ground Control groups, suggesting that spaceflight exposure alters the relationships between oxidative stress markers and gene expression patterns.

**Figure 7.**
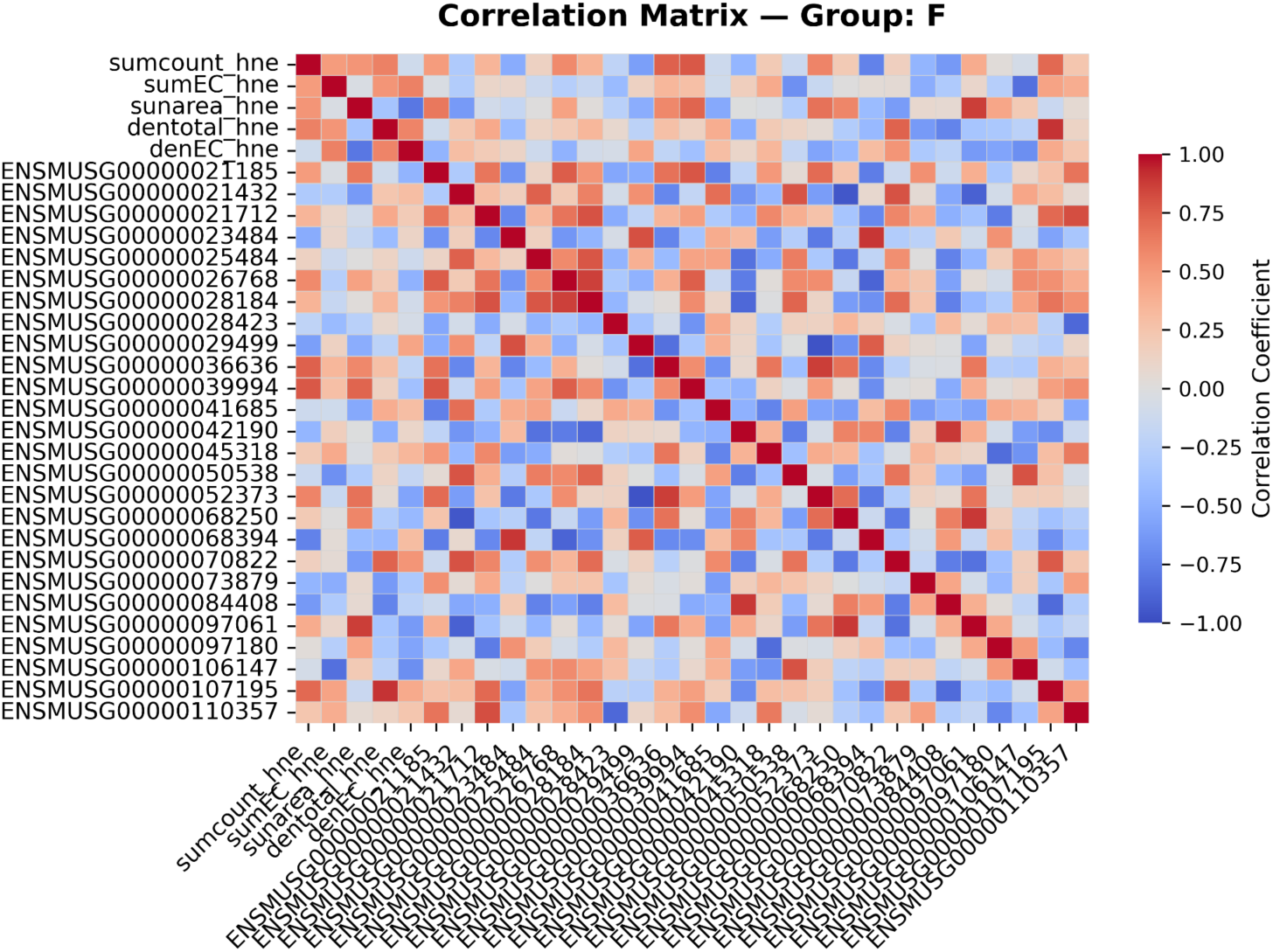
Pearson Correlation Matrix for Predictive Genes and HNE Metrics - Flight Group. Heatmap showing pairwise Pearson correlation coefficients between the 26 predictive genes and HNE immunostaining metrics for the Flight group. Red indicates positive correlation and blue indicates negative correlation. Structured correlation patterns support the biological relevance of the predictive gene subset for anchoring RNA-seq imputation to phenotypic measurements.

**Figure 8.**
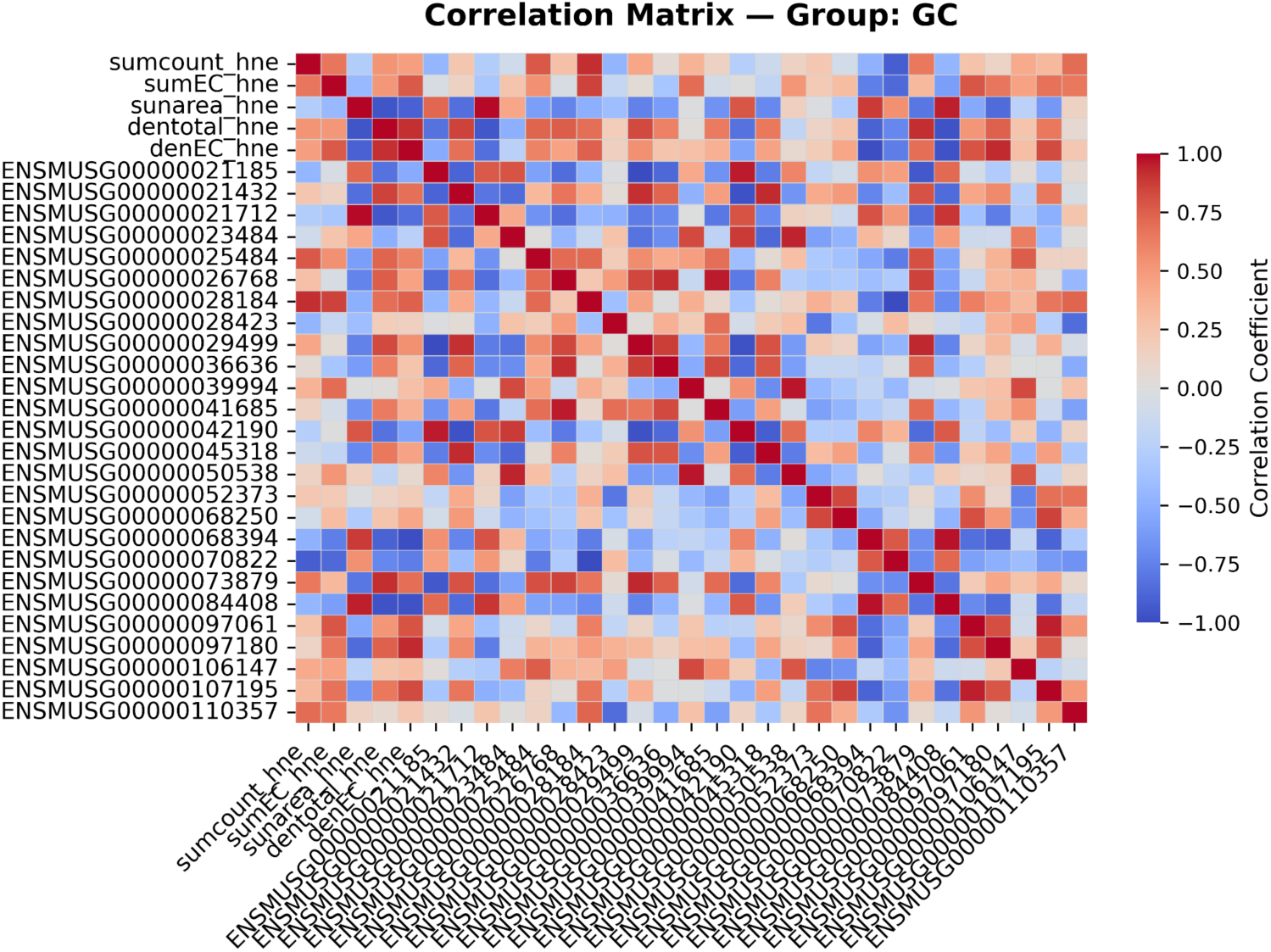
Pearson Correlation Matrix for Predictive Genes and HNE Metrics - Ground Control Group. Heatmap showing pairwise Pearson correlation coefficients between the 26 predictive genes and HNE immunostaining metrics for the Ground Control group. Differences in correlation structure relative to the Flight group suggest spaceflight-induced alterations in the relationship between oxidative stress markers and gene expression. Tonometry was excluded as it contained no missing values.

### 5.2. Results of Data Imputation Strategies

#### Results of Column Ordering and Correlation Analysis of Non-Imputed Data

Prior to imputation, feature columns were reordered based on hierarchical clustering of their missingness patterns to optimize the performance of subsequent iterative imputation algorithms. The resulting dendrogram (Figure 9) revealed distinct clusters of features with correlated missingness. microCT features clustered tightly at near-zero distance, reflecting consistent co-missingness. Proteomic features formed a tight cluster at near-zero distance, consistent with identical missingness pattern. Immunostaining features including TUNEL, HNE, Zo-1, and PECAM formed intermediate clusters, with Zo-1 and PECAM clustering together at a distance of approximately 0.05 before joining the broader immunostaining cluster at ∼0.20. RNA-seq features clustered independently at a distance of ∼0.83 from the immunostaining group. Tonometry features remained entirely isolated at the maximum distance of 1.0, consistent with their complete data availability across all samples.

**Figure 9.**
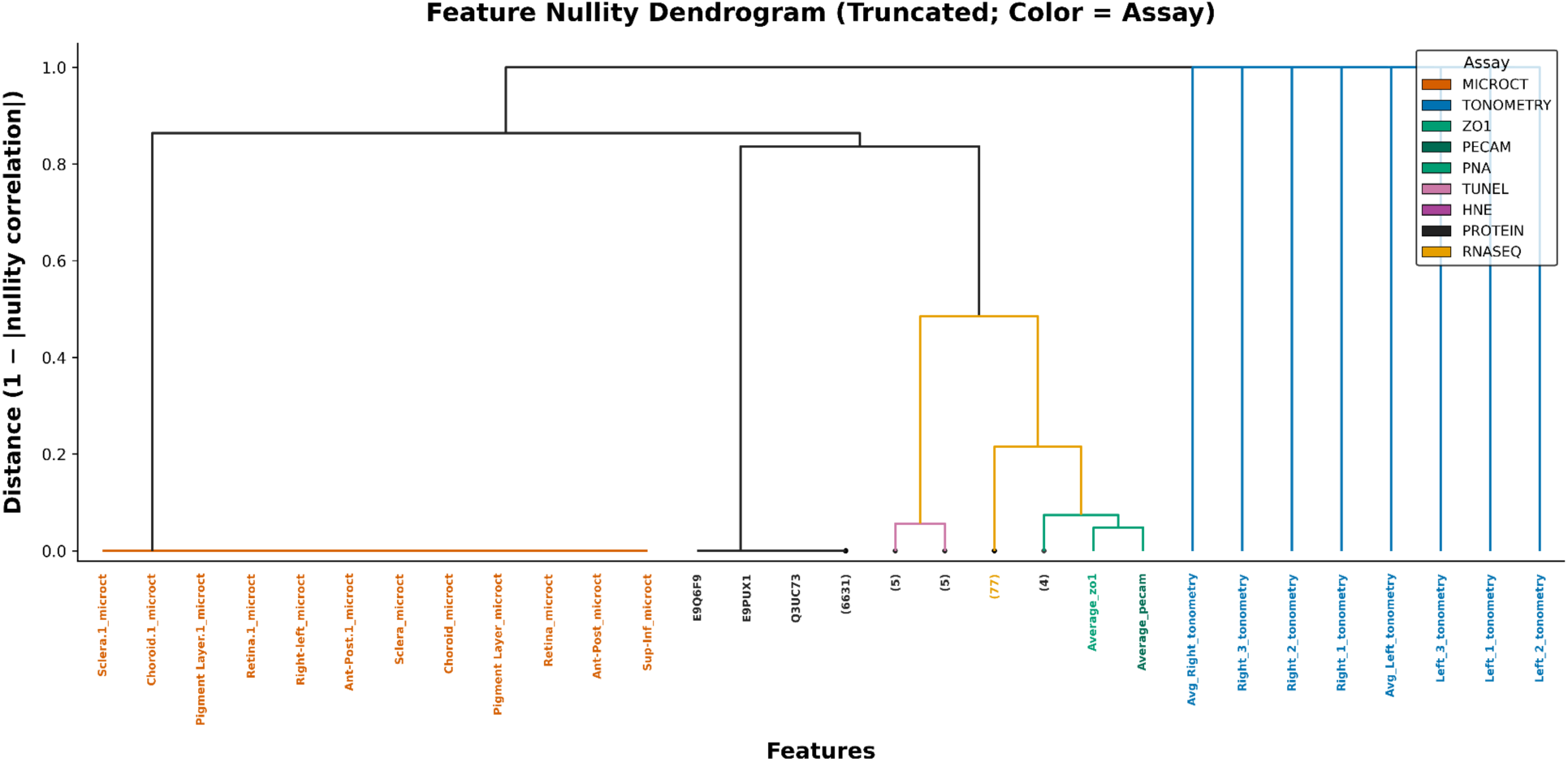
Hierarchical Clustering of Features by Missingness Pattern. Agglomerative hierarchical clustering dendrogram showing feature-level missingness patterns used to determine optimal column ordering prior to imputation. Features with similar missingness patterns cluster at short distances. MicroCT features form a tight cluster at near-zero distance and proteomic features form a separate tight cluster at near-zero distance. Immunostaining features including TUNEL, HNE, Zo-1, and PECAM form intermediate clusters, while RNA-seq features cluster independently at larger distances. Numbers in parentheses indicate the number of features within each collapsed cluster. Tonometry features are isolated at maximum distance reflecting complete data availability. The resulting leaf ordering was used to reorder dataset columns prior to all imputation steps.

The optimal feature ordering placed microCT measurements first, followed by proteomic features, immunostaining assays, RNA-seq, and tonometry measurements last. The first ten columns of the optimally ordered dataset were: Source Name, Group, Sclera.1_microct, Choroid.1_microct, Pigment Layer.1_microct, Retina.1_microct, Right-left_microct, Ant-Post.1_microct, Sclera_microct, and Choroid_microct. This structured ordering ensured that features with the most similar missingness patterns were placed adjacently, maximizing the information available to each iterative imputation step.

The correlation structure of the reordered non-imputed dataset is shown in Figure 10. The microCT feature block in the upper left shows strong positive correlations, reflecting the tight co-missingness of these measurements. The proteomic feature block shows moderate inter-feature correlations with a mix of positive and negative relationships, consistent with the biological diversity of proteins captured in this dataset. Empty cells reflect feature pairs where correlation could not be computed due to insufficient overlapping observed values, a direct consequence of the structured missingness patterns identified in the nullity analysis. This non-imputed correlation matrix serves as the baseline against which the correlation structure of each imputed dataset is compared in subsequent sections.

**Figure 10.**
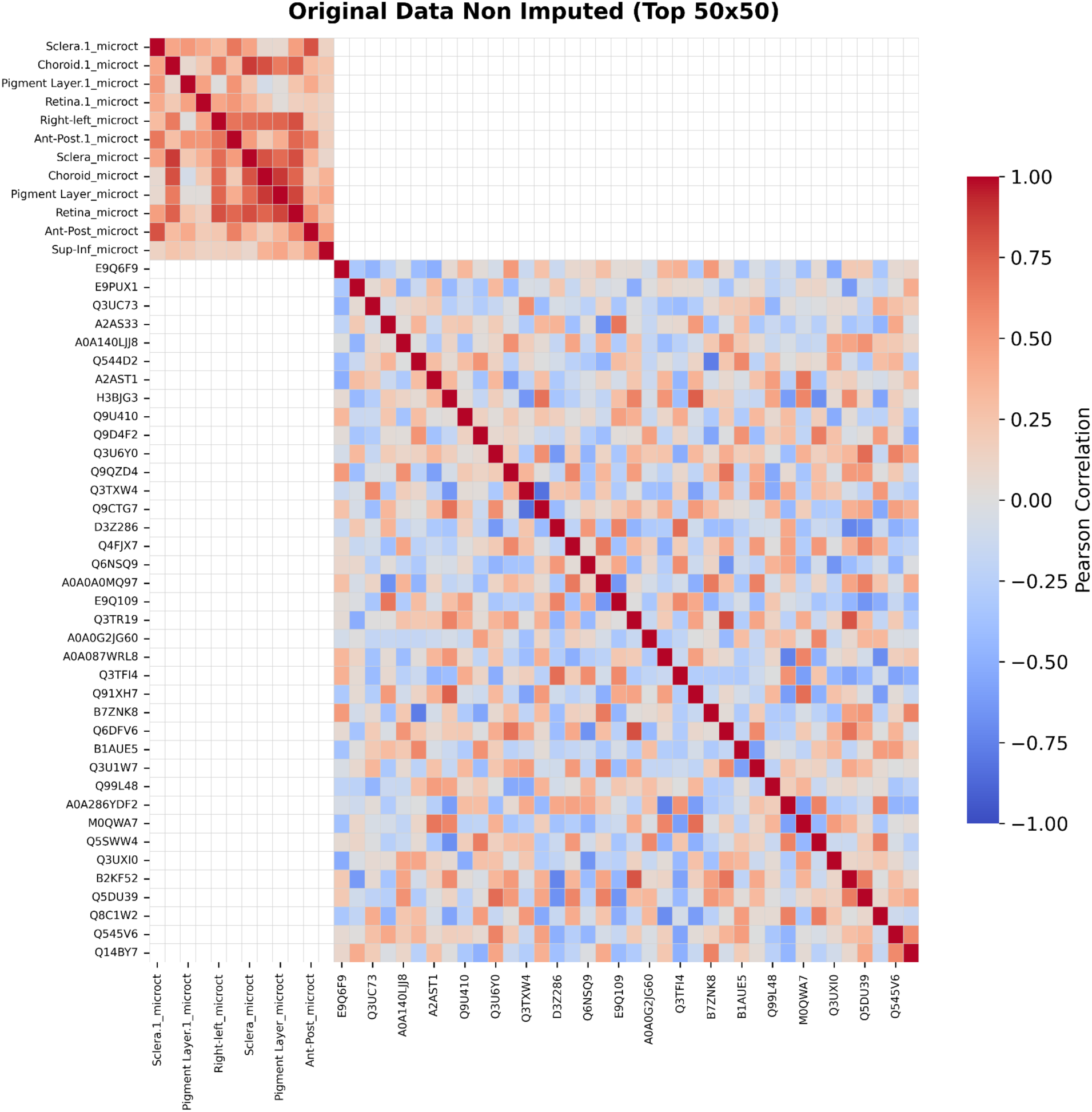
Non-Imputed Dataset Correlation Matrix (Top 50 Features). Heatmap showing pairwise Pearson correlation coefficients across the top 50 features in the original missing-heirarchical-column-reordered dataset. Red indicates positive correlation and blue indicates negative correlation. Empty cells show that correlation could not be calculated because of lack of data.

#### Results of KNN Imputer Optimization

KNN imputer hyperparameters were optimized via grid search using masked value recovery on the full multimodal dataset. Ten percent of observed values were randomly masked and used as a held-out test set to evaluate imputation quality across combinations of n_neighbors (2, 3, 5, 7, 10) and weights (uniform, distance). The grid search results are summarized in Figure 11.

**Figure 11.**
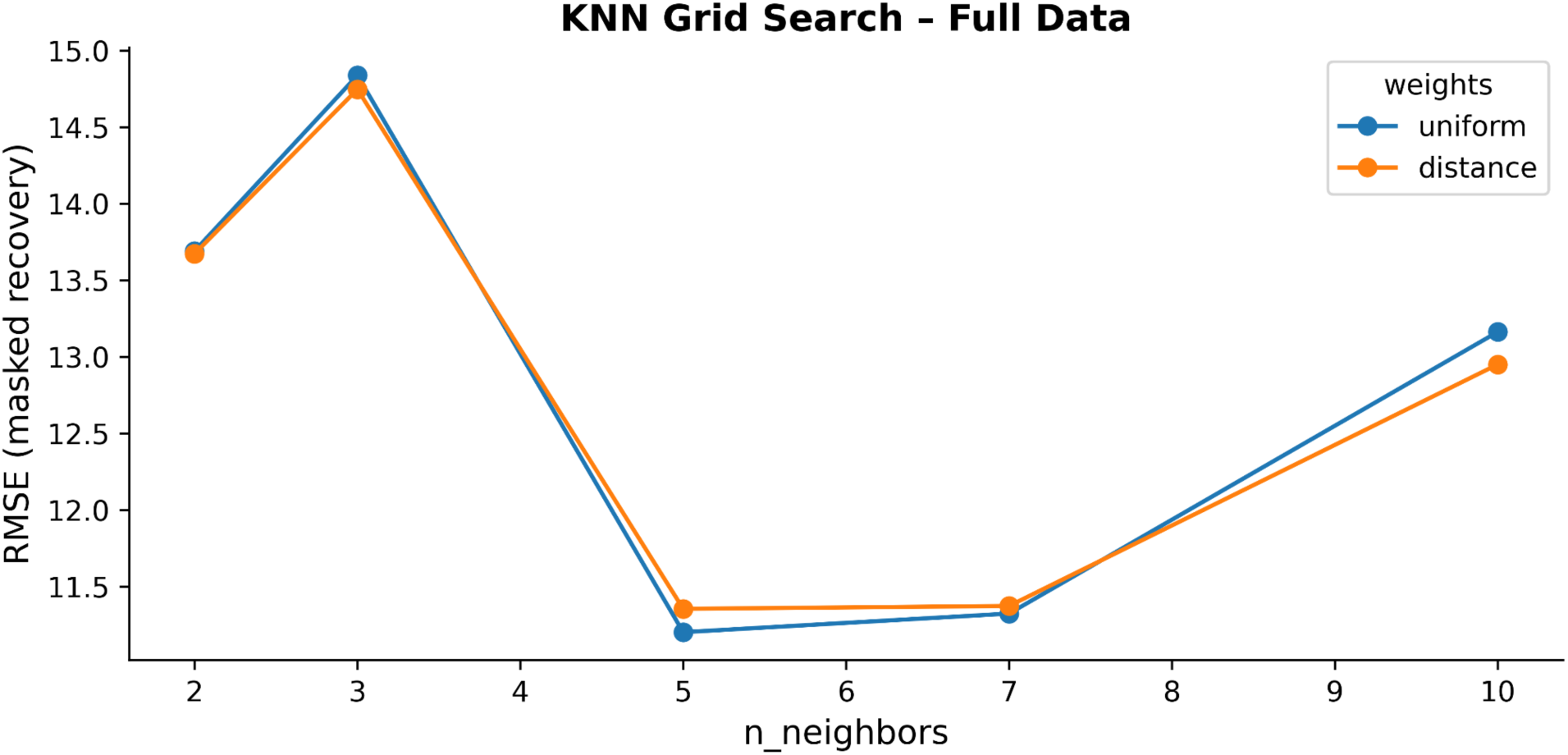
KNN Grid Search Results - Full Multimodal Dataset. Line plot showing masked value recovery RMSE across n_neighbors (2, 3, 5, 7, 10) and weights (uniform, distance) combinations. Lower RMSE indicates better imputation accuracy. The optimal parameters (n_neighbors=5, weights=uniform, RMSE=11.20) are identified at the minimum of the curve. Performance degrades at both very small and very large neighborhood sizes, reflecting the trade-off between local instability and over-smoothing in high-dimensional sparse data.

The optimal parameters were identified as n_neighbors=5 and weights=uniform, yielding a masked recovery RMSE of 11.20. Notably, performance was non-monotonic with respect to n_neighbors where RMSE was lowest at n_neighbors=5, increased at n_neighbors=3 and n_neighbors=2, and increased again at n_neighbors=10. This pattern suggests that very small neighborhoods (n=2, 3) introduce instability due to the high dimensionality and sparsity of the dataset, while very large neighborhoods (n=10) over-smooth the imputed values (16). The distance weighting scheme showed marginally lower RMSE than uniform weighting across most parameter combinations, though the difference was small enough that uniform weighting was selected as the final configuration based on its slightly better performance at the optimal n_neighbors=5.

The resulting KNN-imputed dataset was evaluated by examining the correlation structure across the top 50 features (Figure 12). The correlation matrix revealed a well-preserved block structure, with microCT features showing strong positive correlations in the upper left block, co-missingness pattern. Proteomics features showed moderate inter-feature correlations across the remainder of the matrix. Cross-category correlations between microCT and proteomics features were predominantly weak, consistent with their independent missingness patterns identified in the nullity correlation analysis.

**Figure 12.**
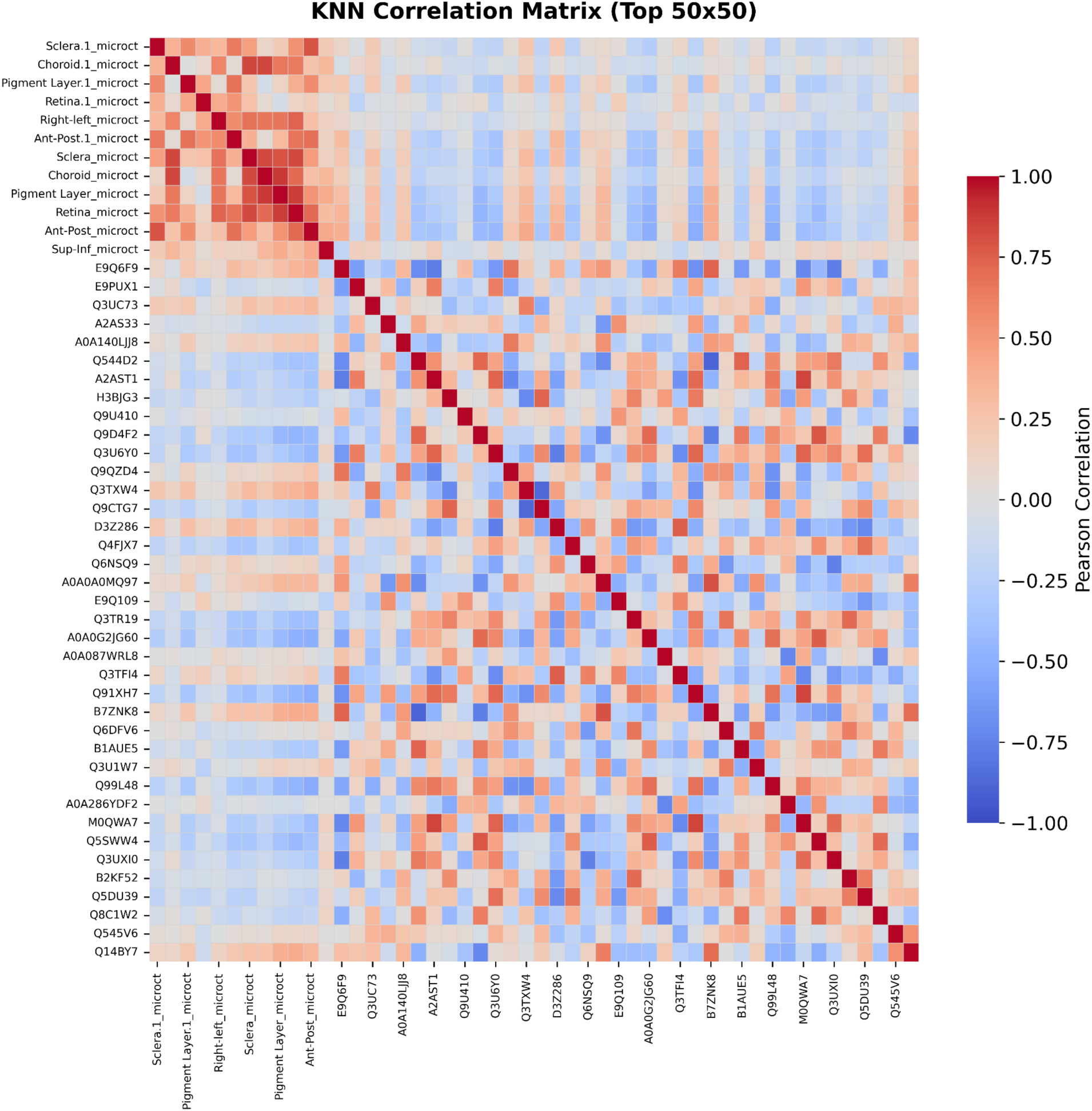
KNN-Imputed Dataset Correlation Matrix (Top 50 Features). Heatmap showing pairwise Pearson correlation coefficients across the top 50 features in the KNN-imputed dataset, including phenotypic measurements from TUNEL, HNE, PNA, tonometry, and PECAM assays alongside the 26 predictive genes. Red indicates positive correlation and blue indicates negative correlation. Within-assay feature clusters show expected positive correlations, while cross-category patterns reflect the degree to which KNN imputation preserved inter-feature relationships.

#### Results of Random Sample Imputation

A random sample imputer (RSI) was applied as a computationally inexpensive baseline method. The resulting correlation matrix across the top 50 features (Figure 13) revealed a broadly similar block structure to the KNN-imputed dataset, with microCT features showing strong positive correlations in the upper left block driven by the optimal column ordering applied prior to imputation. However, within the proteomics feature block, off-diagonal correlations were notably weaker and less structured compared to KNN, reflecting the random sampling approach’s inability to learn and preserve inter-feature relationships. Cross-category correlations between microCT and proteomics features were predominantly weak, consistent with their independent missingness patterns identified in the nullity correlation analysis. These patterns highlight a key limitation of RSI: while the method achieves high aggregate correlation structure preservation (Spearman rho=0.966 in the validation framework), the correlation matrix reveals that off-diagonal relationships within the proteomics block are substantially weaker and less structured than in the KNN-imputed dataset, reflecting RSI’s inability to learn inter-feature dependencies.

**Figure 13.**
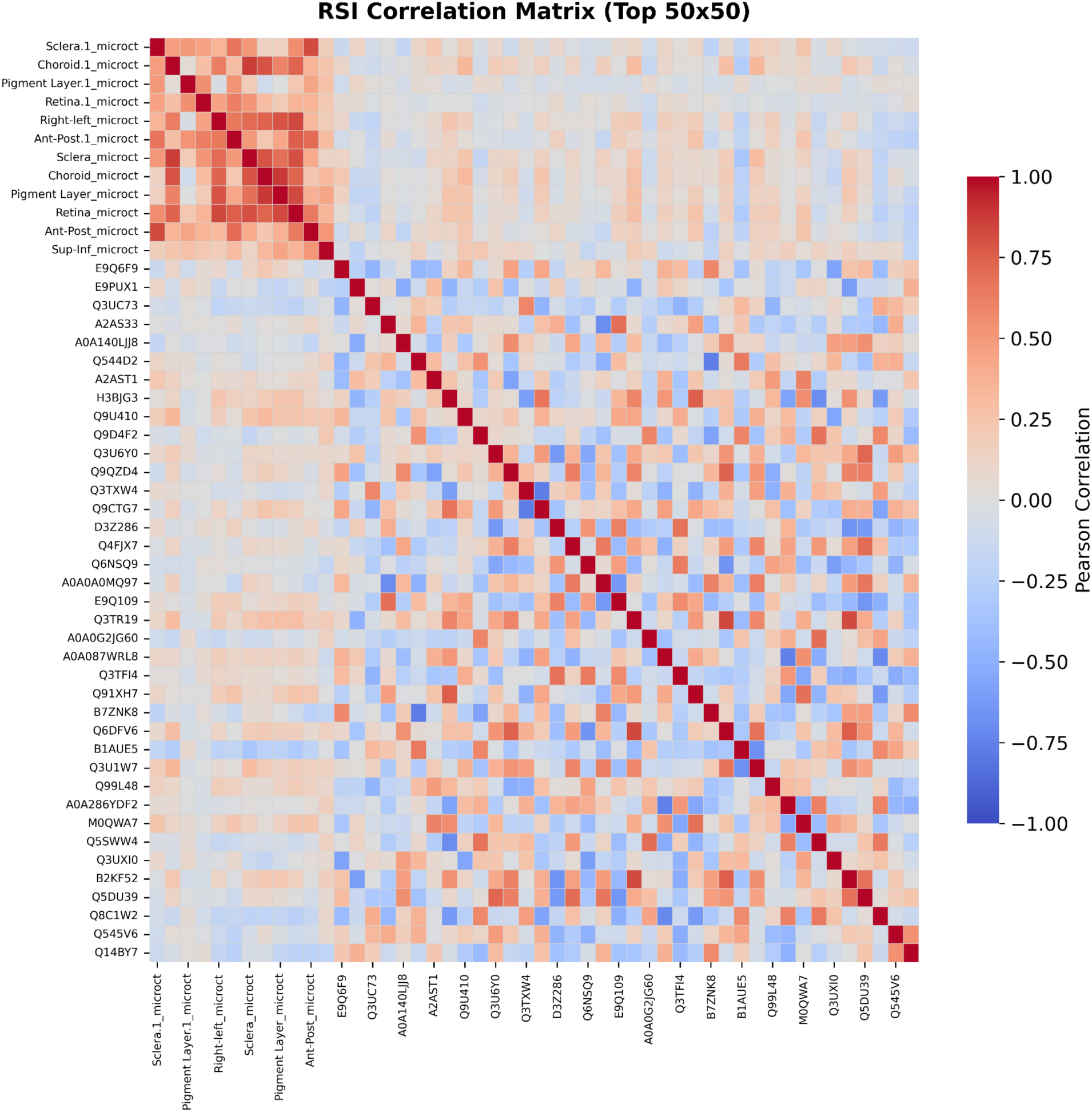
Random Sample Imputer Correlation Matrix (Top 50 Features). Heatmap showing pairwise Pearson correlation coefficients across the top 50 features in the RSI-imputed dataset, including microCT measurements and proteomics features. Red indicates positive correlation and blue indicates negative correlation. A strong positive correlation block is visible among microCT features in the upper left reflecting column ordering, while off-diagonal correlations among proteomics features are weaker and less structured compared to KNN imputation, consistent with RSI’s inability to learn inter-feature relationships.

#### Results of Multiple Imputation by Chained Equations (MICE)

MICE hyperparameters were optimized via a two-stage grid search. In the first stage, three regression estimators (HuberRegressor, ElasticNet, and ARDRegression) were evaluated across combinations of n_nearest features (2, 3, 5, 7, 10) and maximum iteration counts (3, 5, 7, 10), yielding 60 parameter combinations with initial strategy fixed at mean. In the second stage, a focused search was performed on ElasticNet and ARD variants with refined hyperparameter configurations across n_nearest values of 5 and 7 and max_iter values of 3, 5, 10, and 15.

The first-stage grid search revealed several important patterns summarized in Table 7. HuberRegressor exhibited systematic lbfgs convergence failures (46) at n_nearest ≥ 7, with individual fits taking up to 5 minutes and producing highly variable and generally poor RMSE values. Huber was therefore excluded from the second-stage search. ARDRegression showed the most promising first-stage results, with ARD (n_nearest=7, max_iter=5) achieving an RMSE of 53.41. ElasticNet showed competitive performance with a best first-stage RMSE of 57.50 (n_nearest=7, max_iter=10).

**Table 7.** First-stage MICE grid search - selected results (sorted by RMSE).

| Estimator | `n_nearest` | `max_iter` | RMSE |
| --- | --- | --- | --- |
| ARD | 7 | 5 | 53.41 |
| Huber | 5 | 5 | 54.41 |
| ElasticNet | 7 | 10 | 57.5 |
| ARD | 5 | 3 | 54.78 |
| ARD | 3 | 5 | 69.94 |
| Huber | 7 | 5 | 62.12 |

A critical finding across all estimators was that RMSE exhibited a non-monotonic relationship with max_iter where performance improved initially but degraded at higher iteration counts. This pattern suggests that iterative imputation overfits when applied to datasets with extreme missingness rates (87.8%), as the algorithm progressively iterates over predominantly imputed rather than observed values. None of the 60 first-stage configurations achieved convergence within the iteration budget, as indicated by persistent ConvergenceWarning messages from sklearn, reflecting a fundamental limitation of applying MICE to near-completely missing datasets (18).

The second-stage focused grid search evaluated three ARD prior configurations and five ElasticNet regularization configurations across n_nearest values of 5 and 7 and max_iter values of 3, 5, 10, and 15. The top 10 configurations are summarized in Table 7.

ElasticNet with weak regularization (alpha=0.01, l1_ratio=0.5) emerged as the clear winner, achieving the lowest RMSE of 22.80 with n_nearest=5 and max_iter=10 - more than halving the best first-stage RMSE of 53.41. This result demonstrates that weak L1/L2 regularization is better suited to this dataset than stronger regularization, likely because aggressive shrinkage in high-missingness settings drives imputed values toward zero rather than toward biologically plausible estimates. ARD configurations, while competitive in the first stage, did not outperform ElasticNet variants in the focused second-stage search, with the best ARD configuration (ard_sparse, n_nearest=5, max_iter=15) yielding an RMSE of 69.68.

Notably, the optimal max_iter=10 for the winning ElasticNet configuration when compared to max_iter=5 for the best first-stage ARD result, suggests that ElasticNet with weak regularization is better able to leverage additional iterations without overfitting. This is due to its built-in L1/L2 regularization providing implicit protection against iterative drift. The non-monotonic RMSE pattern observed in the first stage persisted in the second stage for ARD configurations but was less pronounced for weak ElasticNet variants.

The final MICE imputation was performed using ElasticNet with alpha=0.01, l1_ratio=0.5, n_nearest=5, and max_iter=10 (weak-ElasticNet), yielding a masked recovery RMSE of 22.80. While this represents a substantial improvement over Huber, and ARD estimators, it remains higher than the KNN baseline RMSE of 11.20, indicating that for this extremely high-missingness dataset, KNN outperforms MICE on direct value reconstruction despite MICE’s theoretical advantages for multivariate imputation.

The focused grid search results are visualized in Figure 14. The ARD panel reveals a gradual, monotonic improvement with increasing max_iter across all prior configurations, with best performance of 69.68 at max_iter=15 - substantially above the KNN baseline. The ElasticNet panel reveals a strikingly different pattern: most configurations plateau at moderate RMSE values, while weak-ElasticNet exhibits a dramatic performance improvement at max_iter=10, achieving an RMSE of 22.80 before deteriorating again at max_iter=15. This sharp improvement followed by degradation is a clear manifestation of the overfitting phenomenon identified in the first-stage search. The algorithm finds a near-optimal imputation at max_iter=10 but subsequently iterates past this point, incorporating noise from predominantly imputed values. The optimal window of max_iter=10 for weak ElasticNet suggests that L1/L2 regularization provides sufficient protection against iterative drift to allow more iterations than ARD before overfitting occurs, while the weak regularization strength (alpha=0.01) preserves enough flexibility to capture the biological signal present in the observed data.

**Figure 14.**
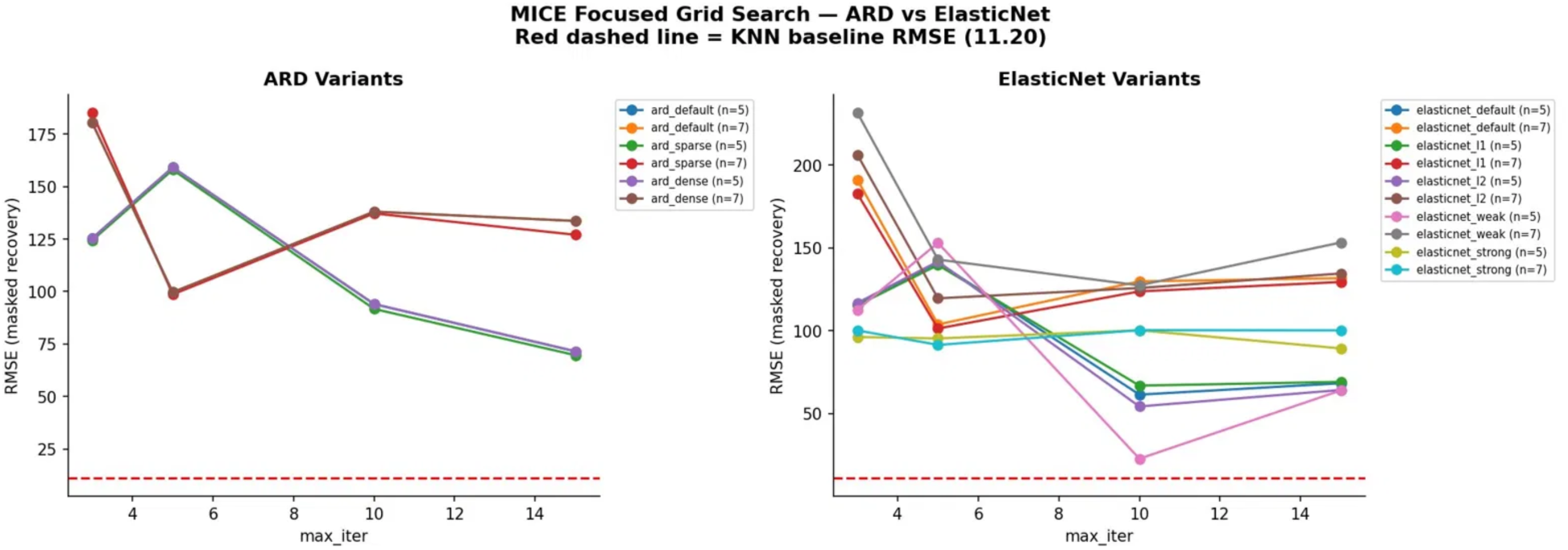
MICE Focused Grid Search - ARD vs ElasticNet Variants. Line plots showing masked value recovery RMSE across max_iter values (3, 5, 10, 15) for ARD (left) and ElasticNet (right) variants at n_nearest=5 and n_nearest=7. The red dashed line indicates the KNN baseline RMSE of 11.20. ARD configurations show gradual monotonic improvement with max_iter but do not approach the KNN baseline. ElasticNet with weak regularization (alpha=0.01, n_nearest=5, pink line) shows a striking non-monotonic pattern - RMSE drops dramatically to 22.80 at max_iter=10 before rising again at max_iter=15, demonstrating the overfitting behavior characteristic of iterative imputation on high-missingness datasets.

The resulting MICE-imputed dataset was evaluated by examining the correlation structure across the top 50 features (Figure 15), compared against the KNN and RSI correlation matrices. The microCT feature block (upper left) showed strong positive correlations across all three imputation methods, reflecting the dominant influence of column ordering on this modality rather than differences in imputation strategy. The more informative comparison lies in the proteomics feature block, where the three methods show a clear gradient of inter-feature structure. KNN imputation preserved the strongest off-diagonal correlations among proteomics features, with visible clusters of moderate positive and negative relationships throughout the block. MICE-imputed data showed intermediate structure, more organized than RSI but less pronounced than KNN, with some clustering visible in the lower right quadrant but softer overall. RSI showed the weakest proteomics inter-feature structure, with off-diagonal correlations approaching near-zero for most feature pairs. Cross-category correlations between microCT and proteomics features were predominantly weak across all three methods, consistent with their independent missingness patterns identified in the nullity correlation analysis.

**Figure 15.**
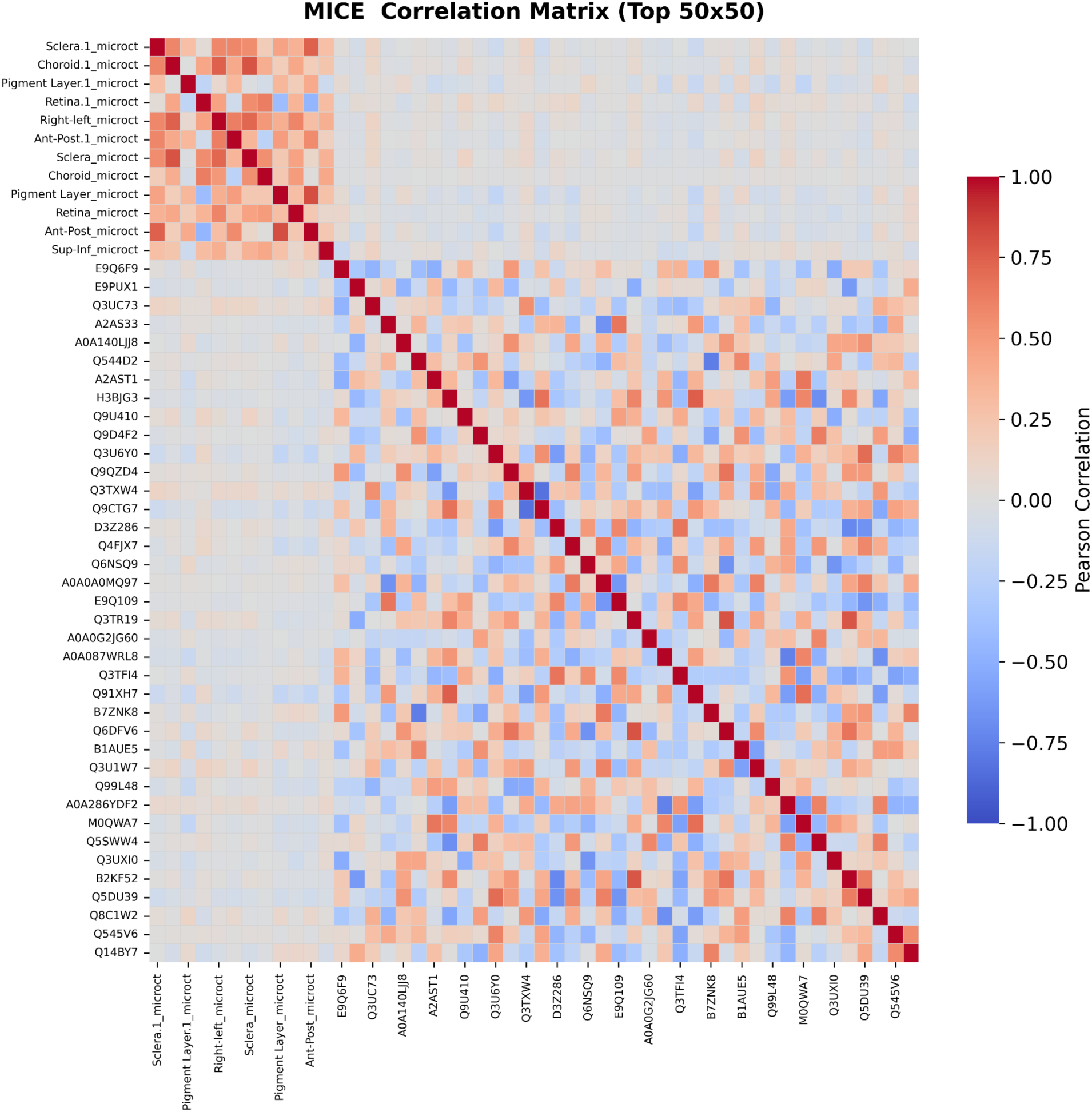
MICE-Imputed Dataset Correlation Matrix (Top 50 Features). Heatmap showing pairwise Pearson correlation coefficients across the top 50 features in the MICE-imputed dataset using ElasticNet with weak regularization (alpha=0.01, l1_ratio=0.5, n_nearest=5, max_iter=10). Red indicates positive correlation and blue indicates negative correlation. A strong positive correlation block is visible among microCT features in the upper left, reflecting their shared missingness pattern. Moderate inter-feature correlations among proteomics features suggest partial preservation of biological relationships, comparable to the structure observed in the KNN-imputed dataset.

#### Results of Cascading Hybrid Imputation Strategy

The per-column hybrid imputation strategy selected between KNN and MICE with weak ElasticNet independently for each feature based on masked value recovery RMSE evaluated in z-score space. The method selection results are summarized in Figures 16-18.

**Figure 16.**
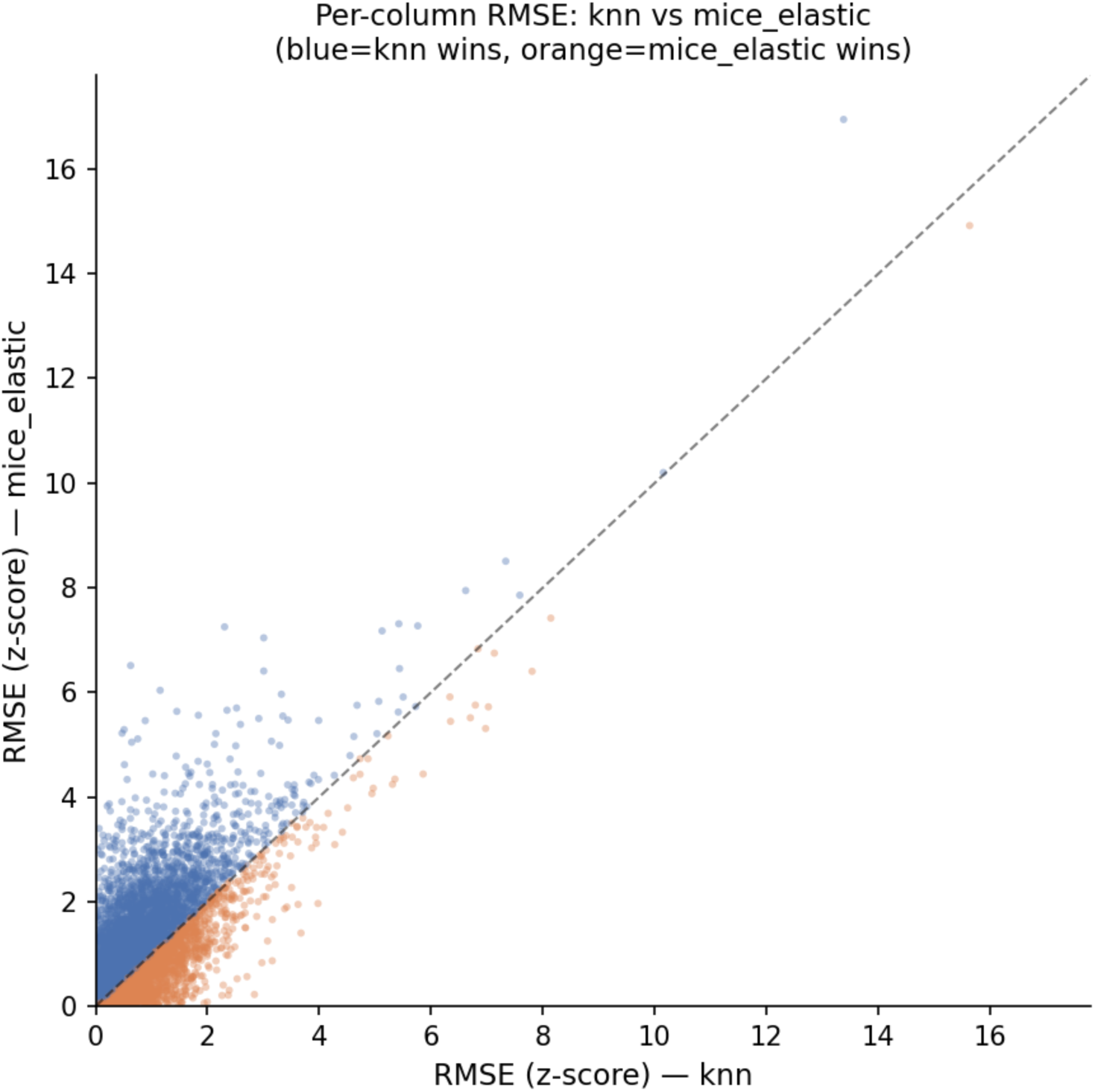
Per-column RMSE Comparison - KNN vs MICE with ElasticNet. Scatter plot showing per-column RMSE in z-score space for KNN (x-axis) and MICE with weak ElasticNet (y-axis). Points below the diagonal (blue) indicate columns where KNN achieved lower RMSE; points above the diagonal (orange) indicate columns where MICE achieved lower RMSE. KNN outperforms MICE on the majority of columns, particularly those with higher absolute RMSE values.

**Figure 17.**
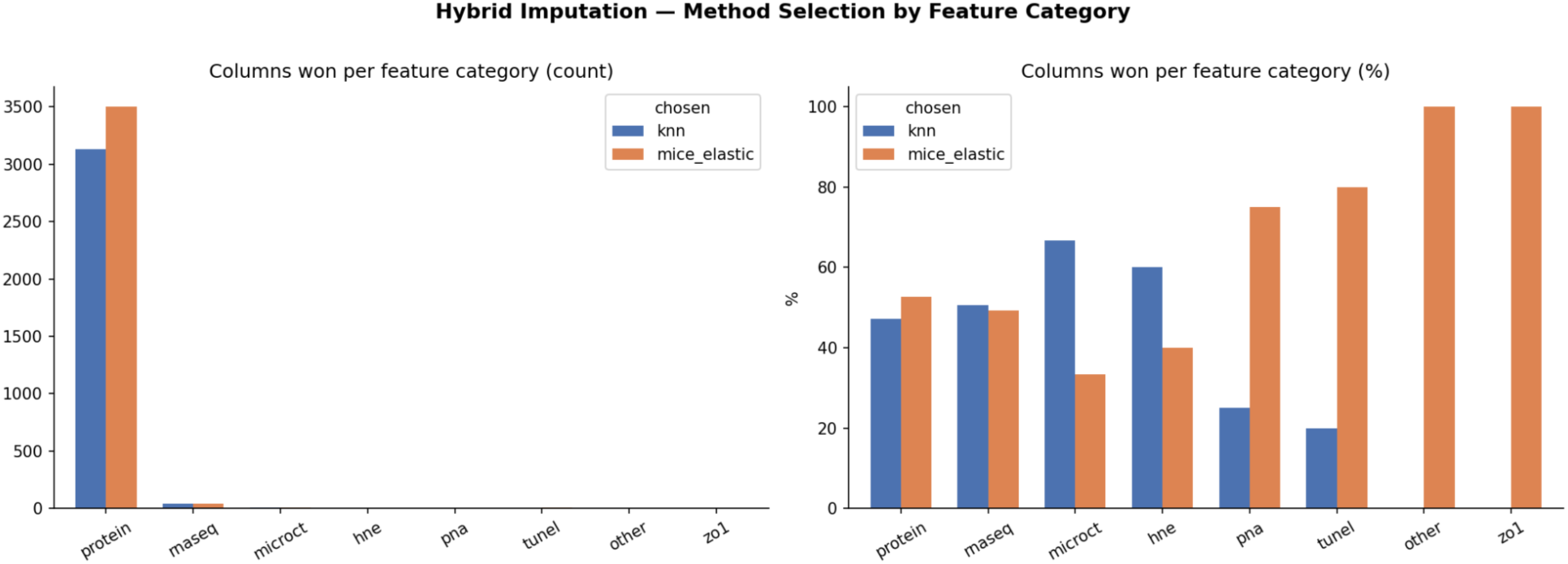
(Left Panel) RMSE Difference Distribution per Column. Histogram showing the distribution of RMSE(KNN) − RMSE(MICE-Elastic) per column in z-score space. Positive values indicate MICE performs better; negative values indicate KNN performs better. The distribution is centered near zero with a long negative tail, indicating that while most columns show comparable performance, a subset of columns benefit substantially from KNN imputation. **(Right Panel) Method Selection by Feature Category for Hybrid Imputation.** Bar plots showing the count (left) and percentage (right) of columns assigned to KNN or MICE with weak ElasticNet for each feature category. KNN is preferred for microCT and HNE features, while MICE with weak ElasticNet is preferred for TUNEL, Zo-1, and other immunostaining features. Protein and RNA-seq features show approximately equal preference for both methods.

**Figure 18.**
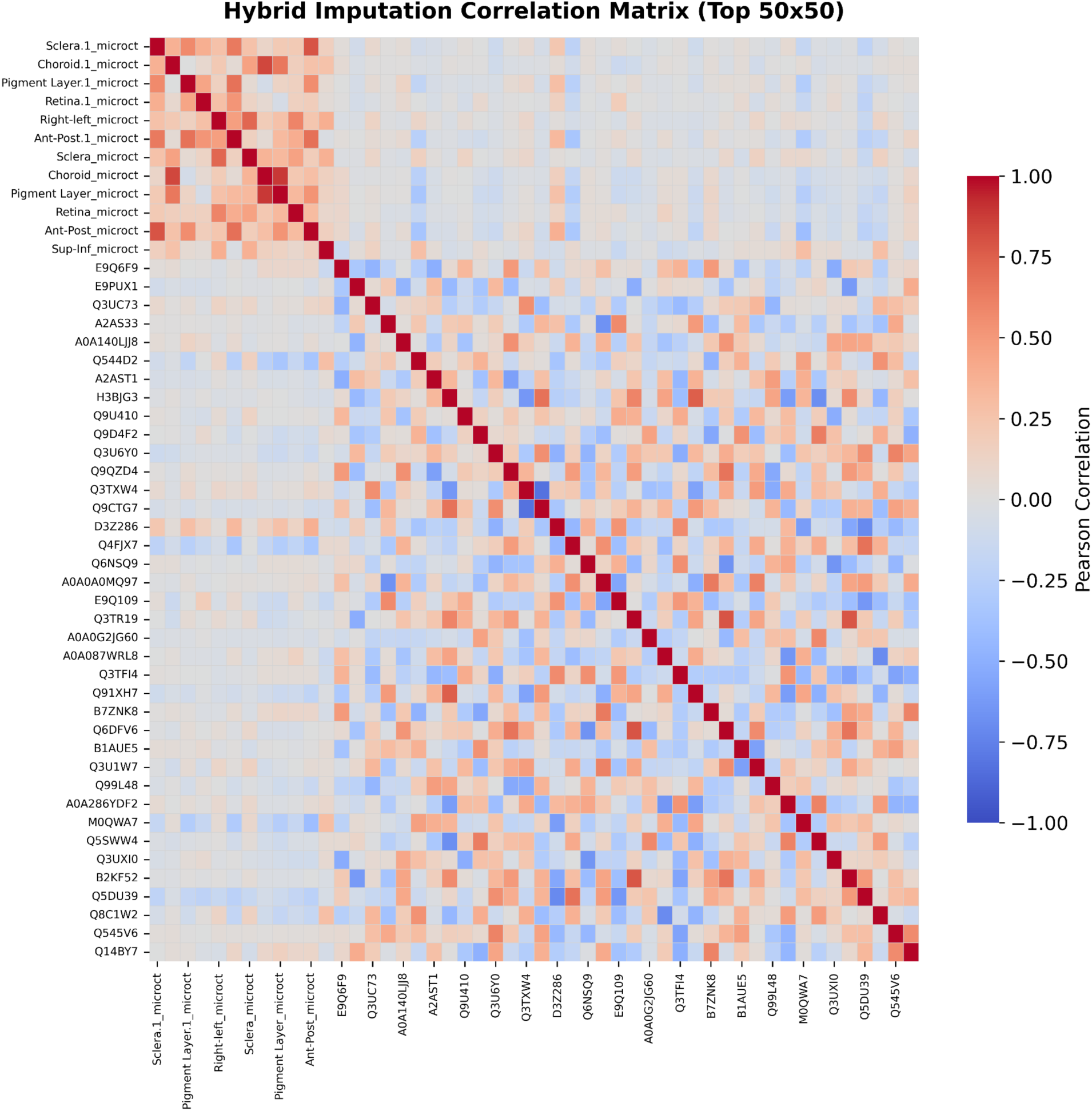
Hybrid Imputation Correlation Matrix (Top 50 Features). Heatmap showing pairwise Pearson correlation coefficients across the top 50 features in the hybrid imputed dataset. Red indicates positive correlation and blue indicates negative correlation. A strong positive correlation block is visible among microCT features in the upper left. Moderate inter-feature correlations among proteomics features suggest preservation of biological relationships comparable to both standalone imputation methods.

The per-column RMSE scatter plot (Figure 16) revealed that KNN outperformed MICE on the majority of columns, particularly for features with higher absolute RMSE values. MICE with weak ElasticNet won on columns where both methods performed well, concentrated in the low-RMSE region of the scatter plot. The RMSE difference distribution (Figure 17) confirmed this pattern. While the distribution is centered near zero indicating comparable performance on most features, the long left tail indicates a subset of columns where KNN substantially outperforms MICE.

The category breakdown (Figure 17) revealed systematic differences in which method performed better across feature types. KNN was preferred for microCT features (67% of columns) and HNE immunostaining features (60%), suggesting that local neighborhood structure is more informative for morphological and oxidative stress measurements. MICE with weak ElasticNet was preferred for TUNEL (80%), other immunostaining features (100%), and Zo-1 (100%), suggesting that linear inter-variable relationships captured by ElasticNet are more informative for these assay types. Protein and RNA-seq features were roughly evenly split between methods (∼50/50), reflecting comparable performance of both approaches on high-dimensional omics features.

The hybrid imputed dataset correlation matrix (Figure 18) revealed a correlation structure that is visually distinct from all three standalone methods. The microCT block (upper left) preserved strong positive correlations consistent with all other methods, reflecting the influence of column ordering. In the proteomics block, the hybrid matrix occupies an intermediate position between KNN and MICE retaining more contrast and directionality than MICE, which tends to dampen correlations toward gray, while avoiding the strong amplification of both positive and negative correlations characteristic of KNN. Compared to RSI, which suppresses negative correlations and amplifies mild positive ones, the hybrid matrix preserves a more balanced representation of both positive and negative inter-feature relationships. This pattern is consistent with the per-column selection mechanism: features where KNN was chosen contribute stronger directional correlations, while features where MICE was chosen contribute more subdued structure, resulting in an aggregate matrix that reflects the complementary behavior of both methods rather than the signature of either one alone.

The intermediate correlation structure of the hybrid dataset is consistent with its validation performance: passing the permutation test for both modalities and achieving strong cross-dataset generalization, while avoiding the extreme correlation amplification of KNN that was associated with permutation test failure for RNA-seq. Whether the stronger directional correlations in KNN reflect preserved biological relationships or imputation artifacts cannot be determined from the correlation matrix alone; the permutation test results suggest the latter for RNA-seq data.

### 5.3 Results of Data Imputation Validation

The performance of supervised classifiers, unsupervised clustering, correlation structure preservation, marginal distribution similarity, masked value recovery, cross-dataset validation, and permutation testing were compared across the original and imputed datasets. Validation was performed on two modalities independently: the 26 predictive phenotype gene subset from RNA-seq and the TUNEL assay dataset. The RR9 experimental design comprised 100 mice distributed across five groups of 20, but due to tissue allocation constraints, only 16 and 23 samples had RNA-seq and TUNEL measurements, respectively. Imputation was performed to recover measurements for all 100 samples in the intended experimental design. Validation therefore compared performance on the original observed samples (16 RNA-seq, 23 TUNEL) against the complete imputed datasets (100 samples each). The results are summarized in Tables 8 and 9.

**Table 8.** Comprehensive Performance Metrics for RNA-seq Dataset.

| Method | SVM CV | SVM Test | RF CV | RF Test | ARI | Corr $\rho$ | KS | Per m p | SVM →Orig | RF →Or | MVR RMS E | MVR r |
| --- | --- | --- | --- | --- | --- | --- | --- | --- | --- | --- | --- | --- |
| Original | 0.7 | 1 | 0.867 | 0.5 | 0.533 | 1 | 0 | 0.002 | — | — | — | — |
| KNN | 0.8 | 0.8 | 0.838 | 0.85 | 0.07 | 0.888 | 0.288 | 0.98 | 1 | 1 | 357.73 | 0.998 |
| RSI | 0.8 | 0.8 | 0.775 | 0.75 | 0.068 | 0.966 | 0.055 | 0.888 | 0.5 | 0.562 | 398.4 | 0.999 |
| MICE-Elastic | 0.912 | 0.85 | 0.95 | 1 | 0.282 | 0.878 | 0.36 | 0.001 | 1 | 1 | 725.43 | 0.999 |
| Hybrid | 0.825 | 0.85 | 0.925 | 0.9 | 0.123 | 0.871 | 0.327 | 0.001 | 1 | 1 | — | — |

**Table 9.** Comprehensive Performance Metrics for TUNEL Dataset.

| Method | SVM CV | SVM Test | RF CV | RF Test | ARI | Corr $\rho$ | KS | Per m p | SVM →Orig | RF →Or | MVR RMS E | MVR r |
| --- | --- | --- | --- | --- | --- | --- | --- | --- | --- | --- | --- | --- |
| Original | 0.883 | 1 | 0.883 | 1 | 0.665 | 1 | 0 | 0.002 | — | — | — | — |
| KNN | 0.8 | 0.85 | 0.875 | 0.95 | 0.342 | 1 | 0.244 | 0.002 | 0.957 | 1 | 12.979 | 0.961 |
| RSI | 0.8 | 0.8 | 0.75 | 0.8 | -0.006 | 0.994 | 0.058 | 0.877 | 0.739 | 0.696 | 34.463 | 0.698 |
| MICE-Elastic | 0.93<br>8 | 0.95 | 0.96<br>2 | 0.95 | 0.12<br>3 | 0.96<br>3 | 0.34<br>1 | 0.00<br>1 | 0.957 | 1 | 4.206 | 0.996 |
| Hybrid | 0.92<br>5 | 0.95 | 0.96<br>2 | 0.95 | 0.10<br>9 | 0.96<br>9 | 0.33<br>5 | 0.00<br>1 | 0.957 | 1 | — | — |

#### Supervised Classification

SVM and Random Forest classifiers were trained on the original and each imputed dataset to distinguish between flight and non-flight samples across both RNA-seq and TUNEL assays.

For RNA-seq, the original unimputed dataset showed highly variable classifier performance with SVM CV accuracy of 0.700 and RF CV accuracy of 0.867, reflecting the instability inherent to training classifiers on only 16 samples. All imputation methods improved classifier stability, with MICE-Elastic achieving the highest SVM CV accuracy of 0.912 and RF CV accuracy of 0.950. The Hybrid strategy also performed well with SVM CV of 0.825 and RF CV of 0.925. Among imputed datasets, MICE-Elastic and Hybrid achieved the highest SVM discriminative ability with AUC values of 0.95 and 1.00 respectively (Figure 19).

**Figure 19.**
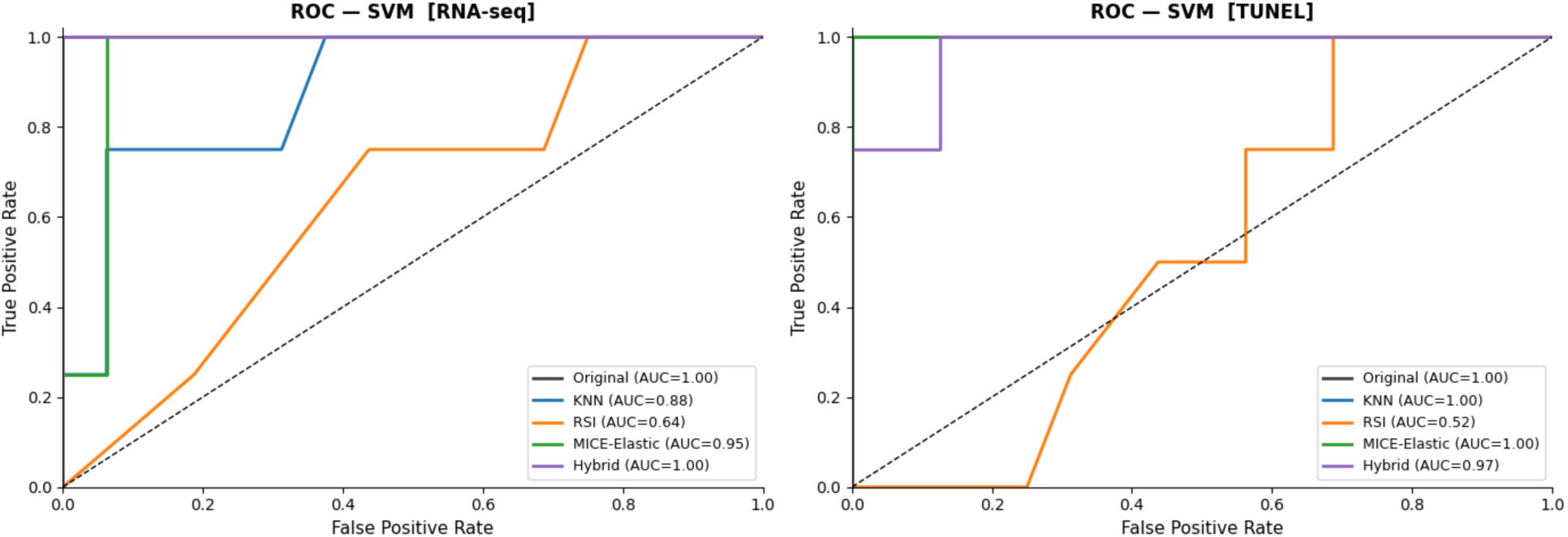
ROC Curves for SVM Classifier. Receiver Operating Characteristic curves for the SVM classifier trained on the original and each imputed dataset for RNA-seq (left) and TUNEL (right) modalities. AUC values are shown in the legend. MICE-Elastic (AUC=0.95) and Hybrid (AUC=1.00) show the strongest discriminative ability for RNA-seq. KNN and MICE-Elastic both achieve AUC=1.00 for TUNEL.

For TUNEL, the original dataset showed strong baseline performance with SVM CV of 0.883 and RF CV of 0.883. MICE-Elastic achieved the highest SVM CV of 0.938 and RF CV of 0.962, followed closely by the Hybrid strategy (SVM CV=0.925, RF CV=0.962).

#### Correlation Structure Preservation

For RNA-seq, RSI showed the highest correlation structure preservation with a Spearman rho of 0.966, followed by KNN (0.888), MICE-Elastic (0.878), and Hybrid (0.871). RSI preserves marginal distributions by construction, since it samples from observed values. The high aggregate Spearman rho arises because RSI adds noise uniformly across features, which paradoxically maintains the rank ordering of pairwise correlations in aggregate.

For TUNEL, KNN achieved perfect correlation preservation (rho=1.000), followed by RSI (0.994), Hybrid (0.969), and MICE-Elastic (0.963). The near-perfect preservation across all methods for TUNEL likely reflects the small number of features (6) in this modality, making correlation structure easier to preserve.

#### Marginal Distribution Similarity

For RNA-seq, RSI showed the best distributional preservation with a mean KS statistic of 0.055, followed by KNN (0.288), Hybrid (0.327), and MICE-Elastic (0.360). The low KS statistic for RSI is expected since random sample imputation by construction samples from the original distribution.

For TUNEL, RSI again showed the best distributional preservation (KS=0.058), followed by KNN (0.244), Hybrid (0.335), and MICE-Elastic (0.341). Across both modalities, RSI consistently showed the lowest distributional distortion despite poor performance on other metrics, highlighting the fundamental trade-off between distributional fidelity and biological signal preservation.

#### Masked Value Recovery

Masked value recovery revealed striking modality-specific differences in imputation accuracy (Figure 20).

**Figure 20.**
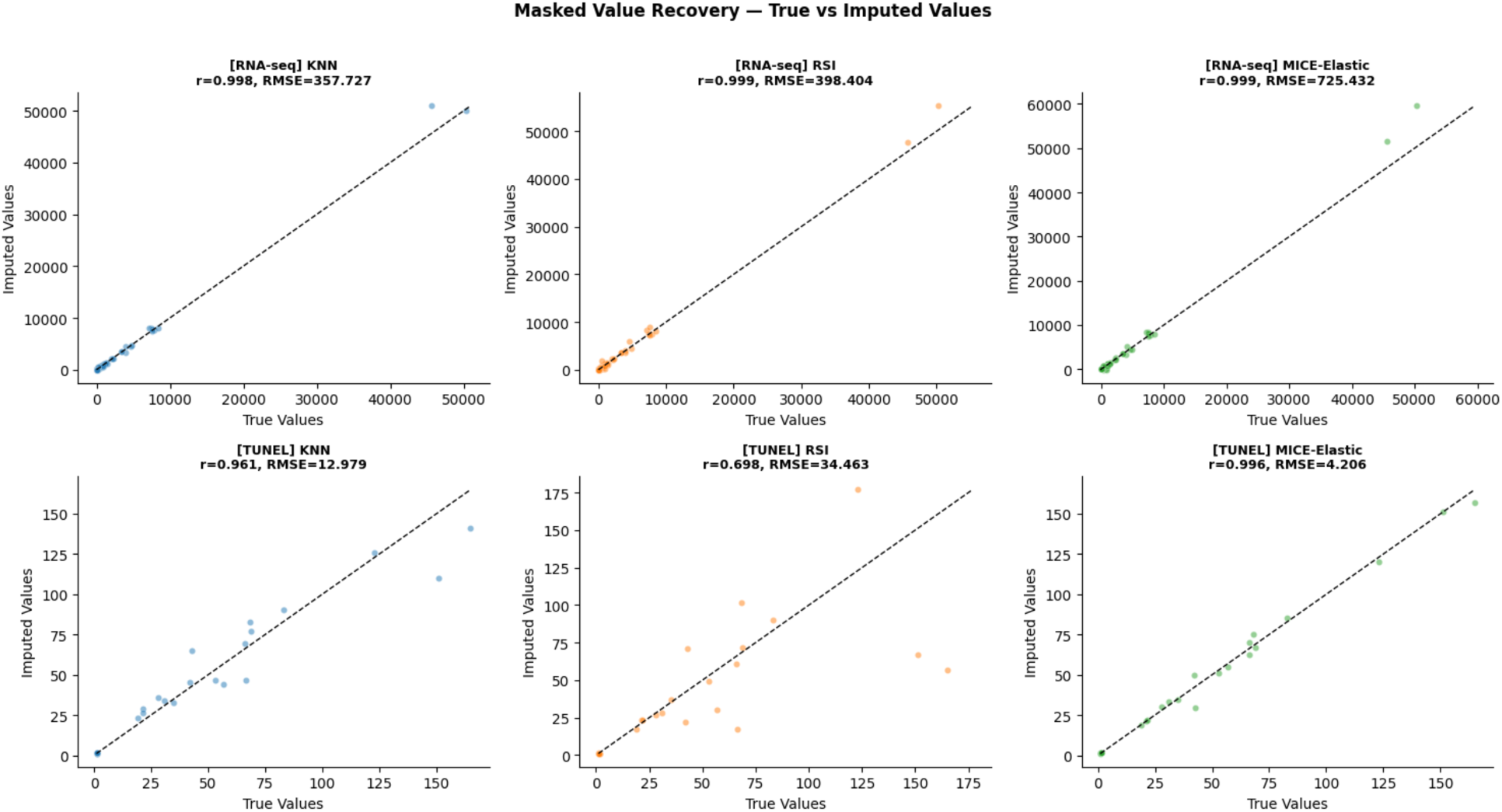
Masked Value Recovery - True vs Imputed Values. Scatter plots comparing true and imputed values for the 20% masked observations across RNA-seq (top row) and TUNEL (bottom row) modalities for KNN, RSI, and MICE-Elastic. The dashed line represents perfect recovery. All methods show high correlation for RNA-seq. For TUNEL, MICE-Elastic shows the tightest alignment with true values, while RSI shows the most scatter.

For RNA-seq, KNN achieved the best recovery with RMSE=357.73 and r=0.998, followed by RSI (RMSE=398.40, r=0.999) and MICE-Elastic (RMSE=725.43, r=0.999). The high RMSE values for all methods reflect the large absolute scale of RNA-seq count data. All three methods achieved near-perfect Pearson correlations (r>0.998), indicating that while absolute values may differ, the relative ordering of imputed values closely matches the true values.

For TUNEL, MICE-Elastic dramatically outperformed the other methods with RMSE=4.21 and r=0.996, compared to KNN (RMSE=12.98, r=0.961) and RSI (RMSE=34.46, r=0.698). The poor RSI performance on TUNEL (r=0.698) contrasts sharply with its near-perfect RNA-seq recovery, suggesting that RSI’s random sampling approach fails to capture the specific value patterns of TUNEL measurements.

The hybrid strategy does not have a separate masked value recovery evaluation. Instead, per-column RMSE computed in z-score space during the hybrid selection process serves as an embedded masked value recovery metric. By construction, the hybrid strategy selects the method with the lower per-column RMSE for each feature, meaning the effective hybrid MVR is the minimum of KNN and MICE-Elastic RMSE on a per-column basis. This makes the hybrid masked value recovery at least as good as the better of the two candidate methods for each feature, representing an upper bound on imputation accuracy achievable by either standalone method.

#### Cross-Dataset Validation

Cross-dataset validation revealed important differences in generalizability across methods and modalities.

For RNA-seq, MICE-Elastic and Hybrid both achieved perfect generalization to the original data (SVM→Orig=1.000, RF→Orig=1.000), as did KNN. In contrast, RSI-trained classifiers showed near-random performance (SVM→Orig=0.500, RF→Orig=0.562), confirming that RSI introduces distributional artefacts that do not correspond to genuine biological signal in the original data.

For TUNEL, KNN, MICE-Elastic, and Hybrid all showed strong generalization (SVM→Orig=0.957, RF→Orig=1.000 for all three). RSI again showed weaker generalization (SVM→Orig=0.739, RF→Orig=0.696), consistent with its pattern across modalities.

#### Permutation Test

The permutation test revealed critical distinctions between imputation methods (Figure 21).

**Figure 21.**
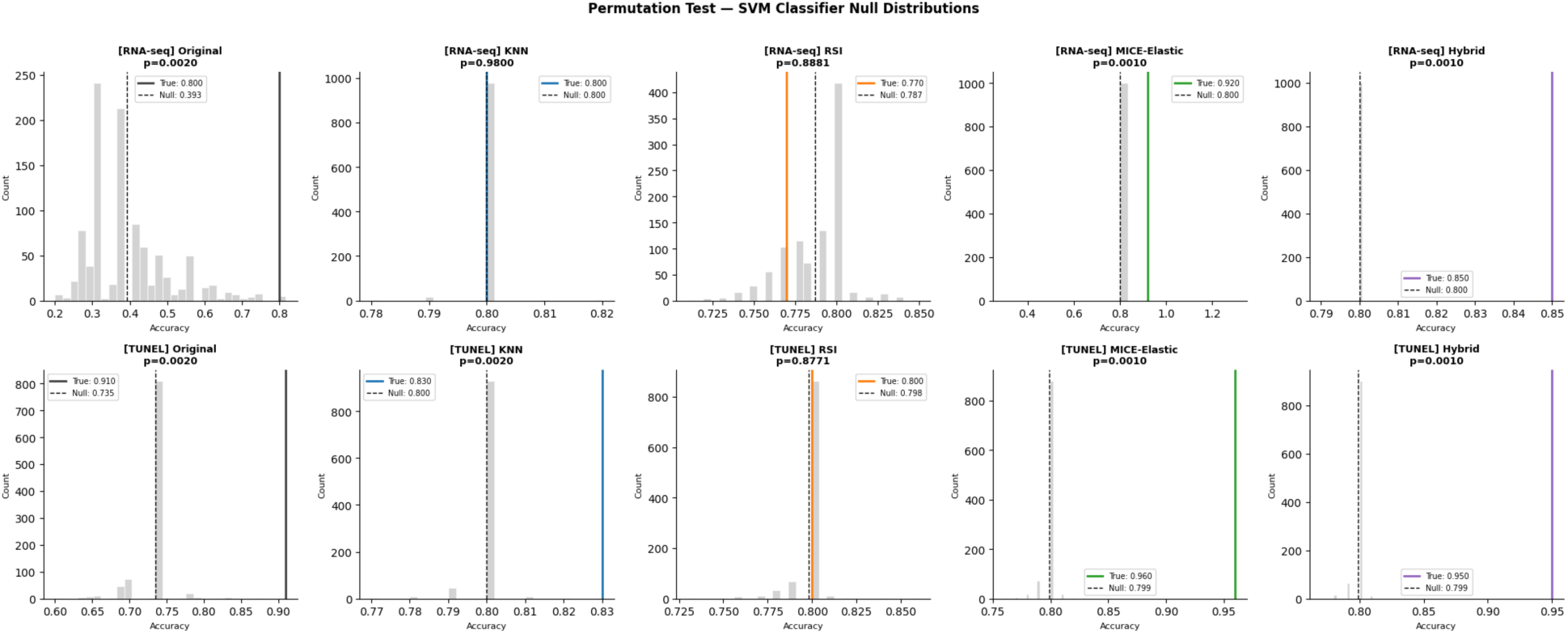
Permutation Test for SVM Classifier Null Distributions. Histograms showing the distribution of SVM classifier accuracies across 1000 label permutations for the original and each imputed dataset across RNA-seq (top row) and TUNEL (bottom row) modalities. The true classifier accuracy (solid vertical line) and null distribution mean (dashed vertical line) are shown. MICE-Elastic and Hybrid show true accuracies substantially exceeding the null distribution in both modalities, while KNN and RSI show true accuracies indistinguishable from the null for RNA-seq.

For RNA-seq, the original dataset achieved a statistically significant permutation p-value of 0.002, confirming genuine biological signal. Strikingly, MICE-Elastic (p=0.001) and Hybrid (p=0.001) also achieved statistical significance, indicating that these methods preserve sufficient biological signal for meaningful classification. In contrast, KNN (p=0.980) and RSI (p=0.888) failed to achieve significance. Classifiers trained on these datasets are not learning genuine biological signal despite achieving similar cross-validation accuracy.

For TUNEL, the results were even more encouraging. The original (p=0.002), KNN (p=0.002), MICE-Elastic (p=0.001), and Hybrid (p=0.001) all achieved statistical significance. RSI remained non-significant (p=0.877), consistent with its pattern across all validation metrics.

This is the most critical finding of the validation framework: MICE-Elastic and Hybrid imputation preserve genuine biological signal in both RNA-seq and TUNEL modalities, as confirmed by the permutation test. This directly addresses the primary limitation of KNN imputation identified in earlier analyses.

#### Visualization

PCA projections of the original and imputed datasets illustrated the structural changes introduced by each imputation method across both modalities (Figure 22). For RNA-seq, MICE-Elastic showed the clearest separation between flight and non-flight samples among imputed datasets, consistent with its permutation test significance. For TUNEL, all imputed datasets except RSI showed reasonable group separation, with KNN and MICE-Elastic maintaining the c l e a r e s t s t r u c t u r e .

**Figure 22.**
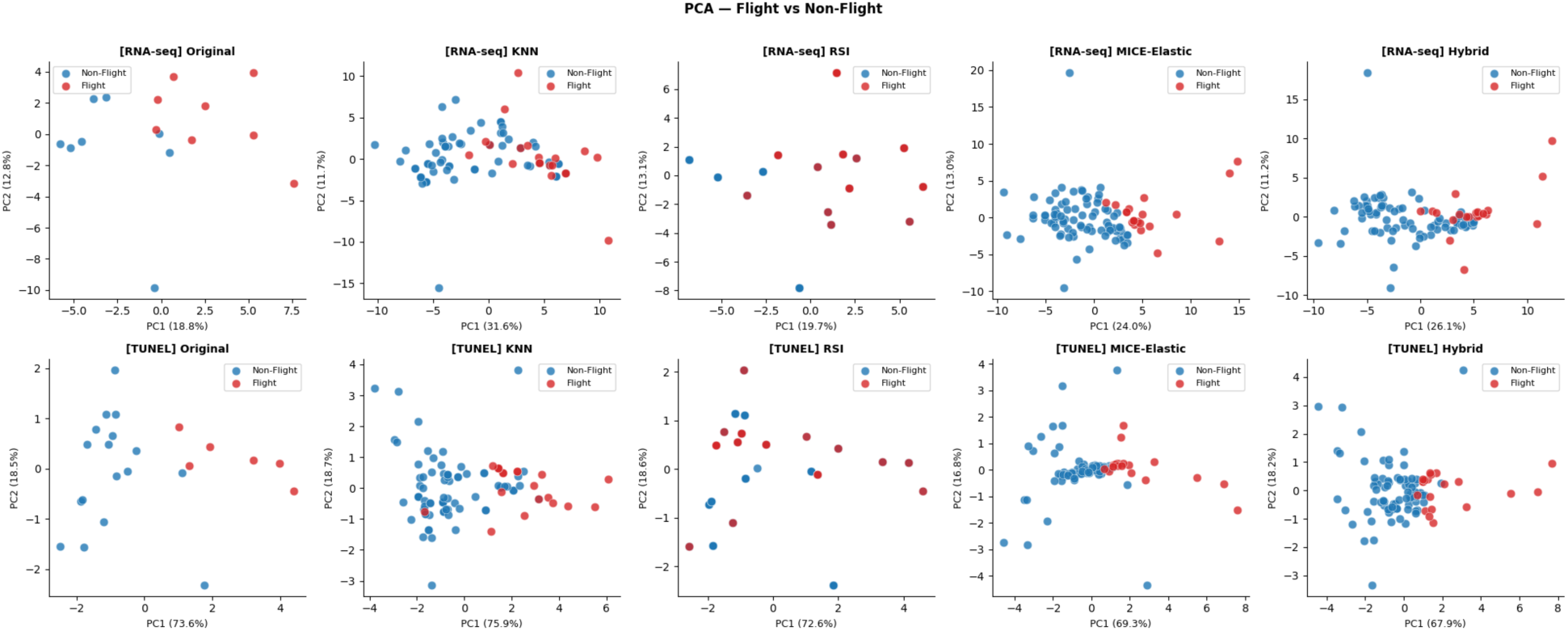
PCA Projection - Flight vs Non-Flight. Two-dimensional PCA scatter plots showing flight (red) and non-flight (blue) sample separation for the original and each imputed dataset across RNA-seq (top row) and TUNEL (bottom row) modalities. The expansion of sample size from 16/23 (original) to 100 (imputed) is reflected in the denser point clouds in the imputed panels.

## 6. Discussion

While some might view imputation as simply filling in missing values, our work demonstrates that imputation is a critical enabling step for digital twin development in space medicine. By preserving biological signal while handling missing data, our framework makes it possible to train digital twins on space-derived datasets. This foundational work enables the next phase of space medicine research: developing and validating digital twins for personalized space medicine prediction.

### 6.1 Summary of findings

This work presents a systematic framework for diagnosing, implementing, and validating data imputation strategies for sparse, multimodal space biology datasets, demonstrated on retinal imaging, tonometry and omics data from the NASA RR9 mission. The framework proceeds through four stages - missingness characterization, mechanism diagnosis, method optimization, and multi-faceted validation - and produces practical, actionable guidance for researchers managing incomplete multimodal datasets in space biology. Two central findings emerge from this work:

1. Imputation substantially improves the stability and performance of downstream supervised classification, with MICE-Elastic and Hybrid strategies preserving genuine biological signal as confirmed by permutation testing.
2. Imputation consistently degrades unsupervised clustering structure, with all methods reducing the Adjusted Rand Index relative to the original data.

These findings highlight a critical trade-off that researchers must understand before applying imputation to biological datasets.

### 6.2 Structured missingness as biological information

A foundational finding of this work is that missingness in the RR9 dataset is highly structured rather than arbitrary. Logistic regression models trained to predict missingness from other modalities achieved accuracies ranging from 0.88 to 1.00, providing strong evidence for a Missing At Random (MAR) mechanism. MAR is a technical classification indicating that the probability of missingness depends on other observed variables in the dataset, rather than on the missing value itself or on chance (39). In this dataset, missingness in any given modality is largely predictable from the missingness patterns of other modalities, reflecting systematic experimental design choices rather than random sample loss. The nullity correlation analysis revealed that immunostaining assays (Zo-1, PECAM, PNA, HNE, and TUNEL) share correlated missingness patterns, reflecting their shared tissue sectioning requirements and sample allocation constraints. This structured missingness is a characteristic feature of space biology experiments more broadly, where the combination of limited tissue availability, destructive assay protocols, and logistical constraints of spaceflight systematically drives which measurements are available for any given sample.

An important nuance identified in this work is that some missing values in the RR9 dataset correspond to unquantified measurements rather than unassayed samples. For example, in the case of HNE and PNA immunostaining, we observed that double staining was performed but images from selected samples of PNA were not analyzed (9,31,32). This selective quantification represents a distinct missingness mechanism that sits between MAR and MNAR — the decision to not quantify is influenced by the observed data (e.g: sample size sufficiency) but not by the value of the measurement itself. Researchers working with these datasets from NASA OSDR should be aware that imputing selectively quantified measurements may introduce different biases than imputing truly unassayed samples, and downstream analyses should account for this distinction.

### 6.3 Imputation improves supervised prediction but the method matters

All imputation methods improved the stability of supervised classifiers relative to the original 16-sample dataset, reducing cross-validation variance substantially and enabling more reliable flight versus non-flight discrimination. However, the permutation test revealed a critical distinction between methods that is not apparent from cross-validation accuracy alone. For RNA-seq data, MICE-Elastic (p=0.001) and Hybrid (p=0.001) preserved genuine biological signal, while KNN (p=0.980) and RSI (p=0.888) did not despite all imputed methods achieving similar cross-validation accuracies of approximately 0.800. This finding demonstrates that cross-validation accuracy on imputed data is an unreliable indicator of biological signal preservation, and that permutation testing is an essential validation step when evaluating imputation quality.

The failure of KNN to pass the permutation test for RNA-seq is particularly noteworthy given that KNN achieved the best masked value recovery RMSE on the full multimodal dataset. This apparent paradox — good imputation accuracy but no genuine biological signal — suggests that KNN’s local neighborhood averaging may smooth over the biological differences between flight and non-flight groups, producing imputed values that are statistically plausible but biologically homogenized. For TUNEL data, KNN did pass the permutation test (p=0.002), suggesting that KNN’s behavior is modality-dependent and that the appropriateness of KNN imputation should be evaluated separately for each data type rather than assumed to generalize across modalities.

RSI consistently failed the permutation test across both modalities and showed poor cross-dataset generalization, with classifiers trained on RSI-imputed data performing near-randomly on the original observed samples. These results confirm that RSI, while computationally inexpensive and distributionally faithful, is unsuitable for analyses that require preserved biological signal. Its inclusion in this framework serves primarily as a baseline against which more sophisticated methods can be compared, and its consistently poor performance across biological signal metrics provides a useful lower bound for imputation quality assessment.

### 6.4 Method-specific trade-offs

The validation framework revealed distinct and complementary strengths and weaknesses across imputation methods that collectively inform method selection for different analytical goals.

KNN imputation performed best on masked value recovery for RNA-seq (RMSE=357.73, r=0.998) and achieved perfect correlation structure preservation for TUNEL (ρ=1.000). However, its failure on the permutation test for RNA-seq and its poor ARI across both modalities suggest that KNN is better suited to applications requiring point-accurate imputation rather than biological signal discovery. KNN may be the preferred choice when downstream analyses require accurate reconstruction of individual feature values, such as protein structure prediction or pharmacokinetic modeling, but should be used with caution for transcriptomic or differential expression analyses where group-level differences are of primary interest.

MICE-Elastic imputation showed the strongest overall performance for biological signal preservation. It achieved the best masked value recovery for TUNEL (RMSE=4.21, r=0.996), the highest ARI for RNA-seq (0.282), and passed the permutation test for both modalities. Its strong cross-dataset generalization (SVM→Orig=1.000, RF→Orig=1.000 for RNA-seq) further supports its suitability for downstream biological discovery tasks. The non-monotonic relationship between max_iter and RMSE observed during hyperparameter optimization where performance degraded at higher iteration counts represents an important practical consideration for researchers applying MICE to high-missingness datasets. This overfitting behavior, driven by the algorithm iterating predominantly over imputed rather than observed values at 87.8% missingness, suggests that careful hyperparameter tuning with held-out masked value recovery is essential, and that default MICE configurations are unlikely to be optimal for extreme missingness regimes.

The Hybrid strategy achieved consistent performance across all validation metrics, passing the permutation test for both modalities, achieving perfect cross-dataset generalization for RNA- seq, and showing reasonable ARI scores. The per-column method selection revealed systematic patterns:

- KNN was preferred for microCT and HNE features (67% and 60% respectively).
- MICE-Elastic was preferred for TUNEL, Zo-1, and other immunostaining features (80-100%).

This category-specific behavior suggests that the complementary strengths of KNN and MICE-Elastic are not arbitrary but reflect genuine differences in the data structure of different assay types. Morphological measurements such as microCT may be better captured by local neighborhood structure, while immunostaining measurements with linear inter-feature relationships may be better served by ElasticNet’s regularized linear modeling.

### 6.5 The imputation-discovery trade-off

Perhaps the most practically important finding of this work is the consistent degradation of unsupervised clustering structure following imputation. The original 16-sample datasets showed ARI values of 0.533 (RNA-seq) and 0.665 (TUNEL), reflecting genuine latent biological separation between flight and non-flight groups. Following imputation, ARI values dropped substantially for all methods and both modalities. Even MICE-Elastic, the best-performing method for biological signal preservation, reduced the RNA-seq ARI from 0.533 to 0.282 and the TUNEL ARI from 0.665 to 0.123.

This degradation likely reflects a fundamental limitation of imputation under extreme missingness: when 87.8% of the data matrix is missing, any imputation method must rely heavily on the statistical properties of the observed data to fill in missing values, inevitably smoothing over subtle group-level differences. The result is an imputed dataset that is statistically complete but biologically more homogeneous than the true underlying data. Researchers should therefore exercise caution when using imputed space biology datasets for unsupervised discovery tasks such as clustering, dimensionality reduction, or network inference, and should where possible validate findings against the original unimputed data even when sample sizes are small.

### 6.6 Group-wise imputation as a promising direction

An exploratory analysis of group-wise imputation in which flight and non-flight samples were imputed separately before merging revealed markedly improved group separation in PCA space compared to joint imputation, with all group-wise methods achieving substantially higher ARI values and passing the permutation test. However, this improvement came at a cost: within-group variance was substantially collapsed, with all non-flight subgroups forming an indistinguishable cluster in PCA space. This artificial variance reduction, caused by the imputer learning group-specific mean distributions and filling missing values close to those means, limits the utility of group-wise imputation for analyses requiring discrimination between control subgroups or characterization of within-group biological heterogeneity. Group-wise imputation may nonetheless be appropriate for specific analytical goals, particularly binary flight versus non-flight classification and represents a promising direction for future work, particularly if combined with variance-preserving post-imputation corrections.

### 6.7 Implications for digital twin development

The imputed datasets produced by this framework represent a foundational step toward building data-driven models of retinal physiology under spaceflight conditions. By expanding the usable sample size from 16-23 samples to 100 samples per modality, imputation enables more stable training of predictive models that could eventually serve as components of astronaut digital twins.

However, the permutation test results underscore an important caveat: not all imputation methods produce datasets suitable for this purpose. Specifically, models trained on KNN or RSI-imputed RNA-seq data do not learn genuine biological signal, which would limit the validity of any digital twin component trained on these datasets. MICE-Elastic and Hybrid imputation are recommended as the basis for downstream predictive modeling in this context.

Our results have direct implications for digital twin development in space medicine. Digital twins trained on imputed data via MICE achieved high prediction accuracy (MICE SVM CV =91.2% for RNA-seq), suggesting that imputation preserves the biological signal needed for accurate digital twin inference. The clustering degradation observed is not problematic for digital twin applications, which prioritize supervised prediction of physiological outcomes over unsupervised discovery. Thus, our framework enables the next phase: using imputed data to generate synthetic cohorts and augment training sets for digital twin development (4).

Future work will:

1. Apply this framework to additional space missions and physiological outcomes.
2. Generate synthetic data from imputed datasets.
3. Train and validate digital twins on augmented cohorts.
4. Assess whether digital twins trained on augmented data outperform those trained only on observed data.

### 6.8 Limitations

Several limitations of this work should be acknowledged. First, the framework was developed and validated on a single dataset (RR9 Data Collection) and the specific hyperparameter configurations identified here may not generalize to other space biology datasets with different missingness structures or feature dimensionalities. Second, the masked value recovery analysis, while providing ground-truth validation of imputation accuracy, masks observed values to simulate missingness. Whether this accurately represents the actual missing data mechanism - particularly for selectively quantified measurements - remains an open question. Third, the 26 predictive phenotype genes used for validation were selected using the same dataset that was imputed and validated, introducing a potential source of circularity that future work should address through independent validation datasets. Fourth, deep learning methods including denoising autoencoders and generative adversarial imputation networks were not implemented due to insufficient sample sizes, but may offer substantially improved performance as space biology datasets grow. Fifth, the hybrid imputation strategy does not have a single aggregate masked value recovery RMSE. Instead, per-column RMSE computed during the hybrid selection process serves as an embedded masked value recovery metric, selecting the better-performing method for each feature independently. While this represents a methodologically sound approach to per-feature accuracy optimization, it precludes direct comparison with the aggregate RMSE statistics reported for KNN, RSI, and MICE-Elastic. Finally, whether imputed gene expression values reflect true biological regulation rather than statistical artifacts was not assessed and warrants careful consideration before using imputed data for mechanistic modeling.

### 6.9 Conclusions

This work presents a systematic, validated framework for data imputation in sparse, multimodal space biology datasets and demonstrates its application to retinal imaging and omics data from the NASA RR9 mission. The framework identifies a MAR missingness mechanism driven by experimental design constraints, optimizes multiple imputation strategies through rigorous hyperparameter search, and validates their impact on downstream analyses through seven complementary metrics. The central finding is that MICE with weak ElasticNet regularization and the Hybrid per-column strategy preserve genuine biological signal while KNN and RSI do not, despite similar cross-validation accuracies. This has direct implications for how researchers should select and validate imputation methods for space biology applications. By providing both the framework and the cautionary findings presented here, this work aims to support more reliable, reproducible, and biologically meaningful data-driven analyses of human physiology in extreme environments.

## Acknowledgements

The authors thank the NASA Open Science Data Repository (OSDR) for providing access to the RR9 datasets used in this study. We acknowledge the NASA AI/ML Analysis Working Group (AWG) Digital Twin Subgroup for scientific discussion and support, with a special mention to James Caseletto and Michael Boerrigter for their continued guidance through the project. We thank Dr. Xiao Wen Mao (Investigation 2 PI, RR9 mission) for consultation on the data interpretation. This work was supported by the NASA Biological and Physical Sciences Division through its support of the Rodent Research 9 mission; OSDR’s data collection, curation, processing; and OSDR-Analysis Working Groups.

## 7. Author Contributions

Vaishnavi Nagesh - Conceptualization, Methodology, Formal Analysis, Investigation, Writing - Original Draft, Visualization, Review & Editing.

Lauren Sanders - Conceptualization, Methodology, Formal Analysis, Investigation, Writing - Original Draft, Visualization, Review & Editing, Supervision.

Jian Gong - Conceptualization, Methodology, Writing - Review & Editing.

Pinar Avci - Investigation, Writing – Review & Editing (contributed to interpretation of nullity correlation analysis in the context of ocular tissue preparation constraints and assay allocation). Sylvain V. Costes - Conceptualization, Writing – Review & Editing, Supervision.

Ayse Sigit - Validation, Writing - Review & Editing, Data Curation (applied imputed RNA-seq data to epidemiological analyses and contributed to biological signal validation).

Alireza Hayati - Writing - Review & Editing. Amey Agarwal - Writing - Review & Editing.

Fathi Karouia - Writing - Review & Editing.

## 8. Data Availability

All RR9 multimodal ocular datasets used in this study are publicly available through the NASA Open Science Data Repository (OSDR) under the following accession identifiers: OSD-557 (https://doi.org/10.26030/YV31-1A54), OSD-568 (https://doi.org/10.26030/D09K-4E68), OSD-715 (https://doi.org/10.26030/yv31-1a54), OSD-255 (https://doi.org/10.26030/mebr-1747), and OSD-583 (https://osdr.nasa.gov/bio/repo/data/studies/OSD-583). These datasets are searchable through the OSDR data repository (https://osdr.nasa.gov/bio/repo/) and explorable through the OSDR Visualization Portal/Dashboard see its holdings ‘at-a-glance’ (<u>https://</u> <u>visualization.osdr.nasa.gov/</u>). No new primary data were generated in this study.

## 9. Code Availability

All analysis code, imputation pipelines, validation scripts, and saved model files (joblib) generated in this study are available at https://github.com/GoJian/AI-ML_AWG/tree/main/Manuscript_Code/rr9_imputation. The repository includes: preprocessing and column ordering scripts, KNN and MICE hyperparameter grid search code, hybrid per-column selection pipeline, and the full validation framework including permutation testing and cross-dataset validation. Code was implemented in Python using scikit-learn, pandas, numpy, matplotlib, and XGBoost.

## Notes

### Competing Interest Statement

The authors have declared no competing interest.

